# Normative Modeling of Brain Morphometry Across the Lifespan Using CentileBrain: Algorithm Benchmarking and Model Optimization

**DOI:** 10.1101/2023.01.30.523509

**Authors:** Ruiyang Ge, Yuetong Yu, Yi Xuan Qi, Yunan Vera Fan, Shiyu Chen, Chuntong Gao, Shalaila S Haas, Amirhossein Modabbernia, Faye New, Ingrid Agartz, Philip Asherson, Rosa Ayesa-Arriola, Nerisa Banaj, Tobias Banaschewski, Sarah Baumeister, Alessandro Bertolino, Dorret I Boomsma, Stefan Borgwardt, Josiane Bourque, Daniel Brandeis, Alan Breier, Henry Brodaty, Rachel M Brouwer, Randy Buckner, Jan K Buitelaar, Dara M Cannon, Xavier Caseras, Simon Cervenka, Patricia J Conrod, Benedicto Crespo-Facorro, Fabrice Crivello, Eveline A Crone, Liewe de Haan, Greig I de Zubicaray, Annabella Di Giorgio, Susanne Erk, Simon E Fisher, Barbara Franke, Thomas Frodl, David C Glahn, Dominik Grotegerd, Oliver Gruber, Patricia Gruner, Raquel E Gur, Ruben C Gur, Ben J Harrison, Sean N Hatton, Ian Hickie, Fleur M Howells, Hilleke E Hulshoff Pol, Chaim Huyser, Terry L Jernigan, Jiyang Jiang, John A Joska, René S Kahn, Andrew J Kalnin, Nicole A Kochan, Sanne Koops, Jonna Kuntsi, Jim Lagopoulos, Luisa Lazaro, Irina S Lebedeva, Christine Lochner, Nicholas G Martin, Bernard Mazoyer, Brenna C McDonald, Colm McDonald, Katie L McMahon, Tomohiro Nakao, Lars Nyberg, Fabrizio Piras, Maria J Portella, Jiang Qiu, Joshua L Roffman, Perminder S Sachdev, Nicole Sanford, Theodore D Satterthwaite, Andrew J Saykin, Gunter Schumann, Carl M Sellgren, Kang Sim, Jordan W Smoller, Jair Soares, Iris E Sommer, Gianfranco Spalletta, Dan J Stein, Christian K Tamnes, Sophia I Thomopolous, Alexander S Tomyshev, Diana Tordesillas-Gutiérrez, Julian N Trollor, Dennis van ’t Ent, Odile A van den Heuvel, Theo GM van Erp, Neeltje EM van Haren, Daniela Vecchio, Dick J Veltman, Henrik Walter, Yang Wang, Bernd Weber, Dongtao Wei, Wei Wen, Lars T Westlye, Lara M Wierenga, Steven CR Williams, Margaret J Wright, Sarah Medland, Mon-Ju Wu, Kevin Yu, Neda Jahanshad, Paul M Thompson, Sophia Frangou

## Abstract

We present an empirically benchmarked framework for sex-specific normative modeling of brain morphometry that can inform about the biological and behavioral significance of deviations from typical age-related neuroanatomical changes and support future study designs. This framework was developed using regional morphometric data from 37,407 healthy individuals (53% female; aged 3–90 years) following a comparative evaluation of eight algorithms and multiple covariate combinations pertaining to image acquisition and quality, parcellation software versions, global neuroimaging measures, and longitudinal stability. The Multivariate Factorial Polynomial Regression (MFPR) emerged as the preferred algorithm optimized using nonlinear polynomials for age and linear effects of global measures as covariates. The MFPR models showed excellent accuracy across the lifespan and within distinct age-bins, and longitudinal stability over a 2-year period. The performance of all MFPR models plateaued at sample sizes exceeding 3,000 study participants. The model and scripts described here are freely available through CentileBrain (https://centilebrain.org/).

## Introduction

Normative modeling is a class of statistical methods to quantify the degree to which an individual-level measure deviates from the pattern observed in a normative reference population. Normative modeling of neuroimaging phenotypes has mostly focused on brain morphometry given the wide availability of structural magnetic resonance imaging (MRI) data^1–4^ with recent extensions into diffusion MRI.^5^ Normative modeling is emerging as a promising new approach to the investigation of brain alternations in neuropsychiatric disorders.^6-11^ However, the value of normative models as research and potentially clinical tools relies on their methodological robustness which has yet to be empirically investigated.

Available normative modeling studies employ a range of linear, nonlinear, and Bayesian algorithms that reflect researchers’ preferences.^1–13^ At present, there is no systematic comparative evaluation of the performance of these algorithms and no empirical determination of the key parameters that may influence model performance. For example, the minimum sample size necessary for reliable normative estimates of brain morphometric measures has not been established and with few exceptions,^1–3,13^ the size of the samples used for the normative reference population is small to modest (range: 145–870).^6–10,14,15^

To address this critical knowledge gap, the aim of the current study is to identify the optimal approach for the normative modeling of brain morphometric data through systematic empirical benchmarking. To this purpose, regional measures of subcortical volume, cortical thickness, and cortical surface area were pooled into a multisite sample of 37,407 healthy individuals (53% female, n=19964, age range: 3–90 years) which was then split into a training and a test subset. Eight algorithms, representing the range of methods currently used in normative modeling studies, were evaluated in terms of their accuracy and computational efficiency, in sex-specific models for each brain morphometric feature. We further quantified the robustness of the different models through extensive empirical testing of the effects of covariates pertaining to image acquisition and parcellation methods, image quality, and global neuroimaging measures, and effects of sample composition (size and age range). The aim was to quantify the accuracy of the different algorithms and identify those parameters that optimise model performance. The results form the basis of CentileBrain (https://centilebrain.org/), an empirically benchmarked framework for normative modeling that is available to the scientific community through a dedicated web platform.

## Methods

### Search strategy and selection criteria

Normative reference values of neuroimaging measures of brain structure and function have great potential as clinical and research tools, but the models used to generate these values must be methodologically robust. We searched electronic databases for articles published in English between Jan 1, 2018, and Jan 31, 2023, using combinations of words or terms that included “normative modeling”, OR “growth curves” OR “centile curves” AND terms referring to specific morphometric features. Although multiple studies employed normative models of brain morphometry, we identified a critical knowledge gap in the paucity of benchmarking statistical methods and sensitivity testing for key parameters that may influence model performance. This study leveraged a large and international sample of healthy individuals (N=37,407) covering the human lifespan to benchmark eight statistical algorithms for normative modeling of brain morphometric measures. In addition to identifying the optimal algorithm, it also defined parameters pertaining to sample composition and size that are essential for robust modeling.

### Samples

We collated de-identified data from 87 datasets from Europe, Australia, USA, South Africa, and East Asia (appendix 1, p 2; appendix 2). Data use aligned with the policies of the ENIGMA Lifespan Working Group (https://enigma.ini.usc.edu/ongoing/enigma-lifespan/), and the policies of individual studies and national repositories. Based on the information provided in each dataset, data were further selected to include high-quality neuroimaging measures (appendix 1, p 3) from participants that were free of psychiatric, medical, and neurological morbidity and cognitive impairment at the time of scanning. Only scans acquired at baseline were included from datasets with multiple scanning assessments. The study design conformed with the STROBE guidelines. Normative models are distinguished into reference models, derived from a sample considered representative of a population in a geographic region at a specific period, and standard models derived from healthy individuals aiming to represent a healthy pattern of age-related changes. Given the nature of our samples, the models developed below are standard models.

### Brain morphometry

Acquisition protocols and scanner vendors varied across datasets (appendix 2). Morphometric feature extraction from whole-brain T_1_-weighted images was implemented using the standard pipelines in the FreeSurfer image analysis suite (http://surfer.nmr.mgh.harvard.edu/) (appendix 2) to yield global measures of total intracranial volume (ICV), mean cortical thickness, and mean surface area, as well as measures of cortical thickness and cortical surface area from the 68 Desikan-Killiany atlas regions (https://surfer.nmr.mgh.harvard.edu/fswiki/CorticalParcellation) and 14 subcortical volumetric measures based on the *Aseg* atlas (https://freesurfer.net/fswiki/SubcorticalSegmentation). Sex-specific normative models were developed separately for each of the 150 regional morphometry measures to accommodate sex differences in brain morphometry.^16^ Sex was determined by self-report. We explored the clustering of the brain morphometry data by geographical regions and did not identify region-specific clusters (appendix 1 p 3).

### Optimization of normative models

The procedures used to generate optimized sex-specific models for each brain morphometric measure are illustrated in figure 1 and consisted of the following steps:

**Figure 1.**
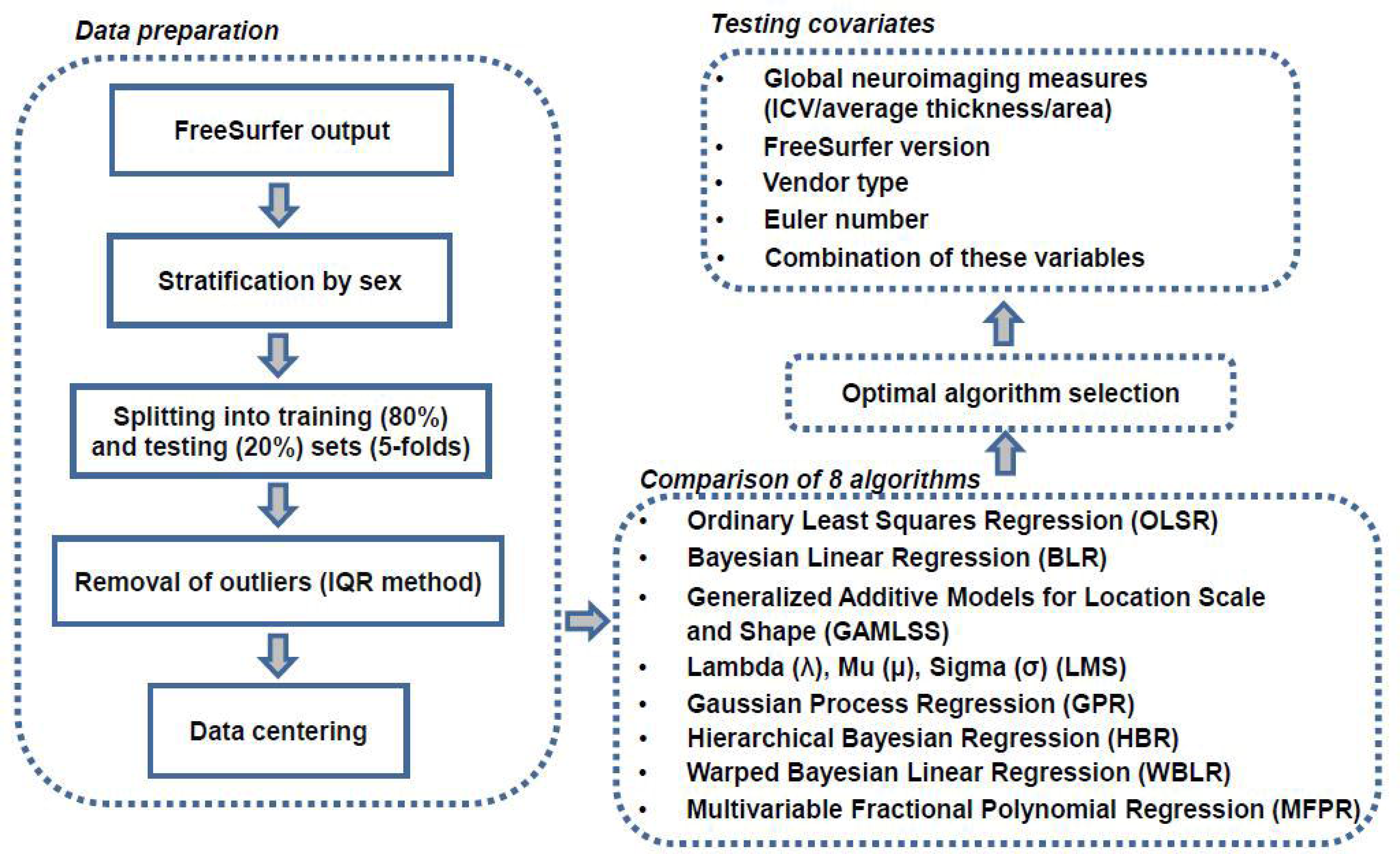
Flowchart of normative model optimization. (1) The study sample was stratified by sex and then split into training (80%) and testing (20%) datasets, followed by outlier removal, and mean-centering; (2) Normative models were generated using eight different algorithms and compared in terms of accuracy and computational efficiency; (3) Explanatory variables were added to identify the appropriate combination for optimal model performance. ICV=intracranial volume; IQR=interquartile range

*(I) Data preparation*: Sex-specific subsamples of the study sample were randomly split into a training subset (80%) and a test subset (20%) stratified by scanning site. Data within the training and testing subset were mean-centered after extreme values, defined as any values greater than 1·5 times the interquartile range (IQR),^17^ in each subset were identified and removed.

*(II) Algorithm selection*: The data for each morphometric measure were analysed with the following algorithms:

(1) Ordinary Least Squares Regression (OLSR) (implemented using the “*lm*” function in R): This is a linear regression model that aims to minimize the sum of squared differences between the observed and predicted values;

(2) Bayesian Linear Regression (BLR) (implemented using the “*stan*” package in R): This is a linear model in which the outcome variable and the model parameters are assumed to be drawn from a probability distribution;

(3) Generalized Additive Models for Location, Scale, and Shape (GAMLSS): This framework can model heteroskedasticity, non-linear effects of variables, and hierarchical structure of the data. This algorithm was implemented using the “*caret*” package in R;

(4) Parametric Lambda (λ), Mu (μ), Sigma (σ) (LMS) method: This subclass of GAMLSS assumes that the outcome variable follows the Box-Cox Cole and Green distribution. This algorithm was implemented using the “*gamlss*” package in R;^15^

(5) Gaussian Process Regression (GPR): This is a nonparametric regression model that follows Bayesian principles, and was implemented using the “*kernlab*” package in R and the “*sigest*” function for estimating the hyperparameter sigma;

(6) Warped Bayesian Linear Regression (WBLR):^18^ This framework is based on Bayesian linear regression with likelihood warping and was implemented using the “PCNtoolkit” (https://github.com/amarquand/PCNtoolkit) in Python following authors’ recommendations;

(7) Hierarchical Bayesian Regression (HBR):^10, 12^ This approach also uses Bayesian principles and is considered particularly useful when variance from multiple hierarchical levels is present, including the scanning protocol or site effects. This algorithm was implemented using the “PCNtoolkit” (https://github.com/amarquand/PCNtoolkit) in Python;

(8) Multivariable Fractional Polynomial Regression (MFPR): This algorithm enables the determination of the functional form of a predictor variable by testing a broad family of shapes and multiple turning points while simultaneously providing a good fit at the extremes of the covariates. This algorithm was implemented using the “*mfp”* package in R and the closed test procedure (known as RA2) was used to select the most appropriate fractional polynomial.

The potential effect of site on performance was addressed both by handling site as a random factor and by site-harmonisation using ComBat-GAM^19^ and then comparing the resulting models.

All models were sex-specific. The total sample of 37,407 healthy individuals comprised 19,964 females and 17,083 males. Each sex-specific sub-sample was divided into the training set (80%) and the testing set (20%) while maintaining the same proportional representation of the sites in the total sample. There was no overlap in terms of participants contributing to the training and the testing sets of each sex-specific subsample. The models were trained using five-fold cross-validation (5F-CV) in the corresponding sex-specific training subset with age being the only explanatory variable. Model parameters were tested in the corresponding sex-specific test subset. In each cross-validation, 80% of the sample was used to train the model and 20% was used to test the model parameters. The mean absolute error (MAE), which is the average of the absolute differences (i.e., errors) between the predicted and the observed data, was averaged across cross-validations and served as the main measure of model performance supplemented by the root mean square error (RMSE), which is the standard deviation of the prediction errors, which was also averaged across cross-validations and by Explained Variance (EV). The computational efficiency of each model was assessed using the central processing unit (CPU) time of the supercomputing infrastructure of the Icahn School of Medicine at Mount Sinai (https://labs.icahn.mssm.edu/minervalab/resources/hardware-technical-specs/).

*(III) Model optimization*: Model optimization involved the evaluation of improvements in the MAE (and RMSE and EV) by adding the following explanatory variables: (1) global neuroimaging measures (i.e., ICV, mean cortical thickness or mean cortical surface area, as appropriate); both linear and non-linear contributions from these variables were considered; (2) scanner vendor type; (3) FreeSurfer version; (4) Euler Number (EN) for scan quality, and (5) combinations of these variables. Each model was trained using 5F-CV in the corresponding sex-specific training subset and then tested in the corresponding sex-specific test subset. Variables that significantly improved performance were retained.

Across regional morphometric measures (and separately in males and females), the MAEs and RMSEs of the optimised models generated by each algorithm were concatenated as a single vector to enable pairwise comparisons between algorithms. Results were considered significant across comparisons at P_FDR_<0·05 using false discovery rate (FDR) correction for multiple testing. Upon completion of steps (*I)–(III),* optimised sex-specific and region-specific models were defined based on the best-performing algorithm and covariate combination. The normative deviation score for each region ^4, 11^ was defined as: 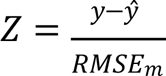 where *y*^^^ is the predicted value, *y* is the observed value and RMSE_m_ is the value in the pre-trained model.

### Sensitivity analyses

*Sample size*: The study sample was partitioned into 75 sex-specific random subsets comprising between 200 to 15,000 participants in increments of 200. The robustness of the optimised sex- and region-specific models to sample size in terms of MAE and RMSE was assessed in each partition using 5F-CV.

*Distinct age bins:* Model accuracy may be influenced by the sample’s age range and by distinct challenges encountered in scanning different age groups such as higher levels of motion in pediatric populations.^20^ Accordingly, the study sample was divided into nine sex-specific age-bins (i.e., age≤10 years; 10<age≤20 years; 20<age≤30 years; 30<age≤40 years; 40<age≤50 years; 50<age≤60 years; 60<age≤70 years; 70<age≤80 years; 80<age≤90 years). The MAE and RMSE of each optimised sex- and region-specific model were estimated in each age bin using 5F-CV. Subsequently, Pearson’s correlation coefficients were computed between the MAE and RMSE values of the models within each sex-specific age bin with those derived from the sex-specific subset of the entire sample. Before computing Pearson’s correlation coefficients, the assumption of linearity was verified by using the Kolmogorov–Smirnov tests and illustrated in scatter plots between the MAE and RMSE values of the models within each sex-specific age bin and those derived from the sex-specific subset of the entire sample (appendix 1 p 16).

*GAMLSS:* As the GAMLSS algorithm is particularly popular for normative modeling, ^21^ we performed additional sensitivity analyses for different GAMLSS models and software packages as detailed in appendix 1, pp 15–19.

*Optimised model performance in longitudinal datasets:* The Southwest Longitudinal Imaging Multimodal Study (SLIM) (http://fcon_1000.projects.nitrc.org/indi/retro/southwestuni_qiu_index.html) and the Queensland Twin Adolescent Brain Study (QTAB) (https://openneuro.org/datasets/ds004146/versions/1.0.4) were used to test the longitudinal stability of the optimal normative models. There is no participant overlap between the SLIM and QTAB and between either dataset and the sample used for model development. The SLIM dataset includes 118 healthy individuals (59 females/59 males; sample age range: 17–22 years for the baseline scans and 19–25 years at follow-up scans) who were rescanned with a mean interval of 2·35 years. The QTAB dataset includes 259 healthy individuals (129 females/130 males; sample age range 9–14 years for the baseline scans and 10–16 years at follow-up scans) who were rescanned with a mean interval of 1·76 years.

### Relevance of normative models of brain morphometry for mental illness

We tested whether normative brain regional Z-scores have an advantage over the observed morphometric measures in predicting diagnostic status and symptom severity using psychosis as an exemplar. To achieve this, we downloaded and parcellated (using FreeSurfer version 7.1.0) T_1_-weighted images from the repository of the Human Connectome Project-Early Psychosis Study (HCP-EP; https://www.humanconnectome.org/study/human-connectome-project-for-early-psychosis). The HCP-EP cohort comprises 91 individuals with early psychosis and 57 healthy individuals (total sample: 48 females/100 males; age range 16·67–35·67 years).^22^ Then each of the algorithms examined here was applied to generate brain regional Z-scores in the HCP-EP cohort.

For diagnostic status prediction, and for each algorithm, the regional Z-scores and the observed neuromorphometric data were entered into separate support vector classification (SVC) models with a linear kernel from the scikit-learn package (version 1.2.2) following established procedures.^23^ The area under the receiver operating characteristic curve (AUC), averaged across all folds within a 5F-CV framework repeated 100 times, was used to evaluate the classification accuracy of each SVC model. Statistical significance was established by comparing the averaged AUC of each model to a null distribution generated from a model trained on 1,000 random permutations of the diagnostic labels (i.e., patient or healthy individual in the HCP-EP cohort). To compare the classification accuracy of the SVC models using the regional Z-scores with the SVC model using the observed neuromorphometric data, we calculated pairwise Δ_*AUC*_ and we tested whether they exceeded chance probability compared to a null distribution using permutation.

For the prediction of symptom severity, and for each algorithm, regional Z-scores and the observed neuromorphometric data were entered into separate ridge regression models with 100 repeats of 5F-CV to predict the psychosis score of the Positive and Negative Syndrome Scale (PANSS) ^24^ in the HCP-EP patients. The MAE of each model, averaged across folds, was used as the performance metric. Within each fold, we applied principal component analysis to reduce the dimensionality of the brain regional measures to the first ten principal components that explained at least 90% of the variance. To compare the predictive accuracy of the regression models using the Z-scores to the model using the observed neuromorphometric data we calculated pairwise Δ_*MAE*_ and followed the same procedures as for the classification models. In the case of predictive accuracy, permutations involved shuffling the PANSS scores of the HCP-EP cohort.

## Results

*Identification of the best-performing algorithm*: In figure 2, the MAE, RMSE, EV and CPU time of the models for the left thalamic volume and left medial orbitofrontal cortical thickness and surface area in females are shown as exemplars (the corresponding data for males in appendix 1, p 4, figure S3). The pattern was the same for all regions across sex-specific models as shown in appendix 1 (pp 5–6; figures S4–S5) and appendix 3. Across all models, the OLSR and MFPR had the shortest CPU times (less than one second) while GPR had the longest (25–60 minutes). Across all sex- and region-specific models, the LMS, GPR, WBLR, and MFPR had comparable values for MAE, RMSE, and EV that were statistically better at P_FDR_<0·05 than those for GAMLSS, BLR, OLSR, and HBR. Accordingly, the MFPR emerged as the preferred algorithm given its combined advantages in accuracy and computational efficiency.

**Figure 2.**
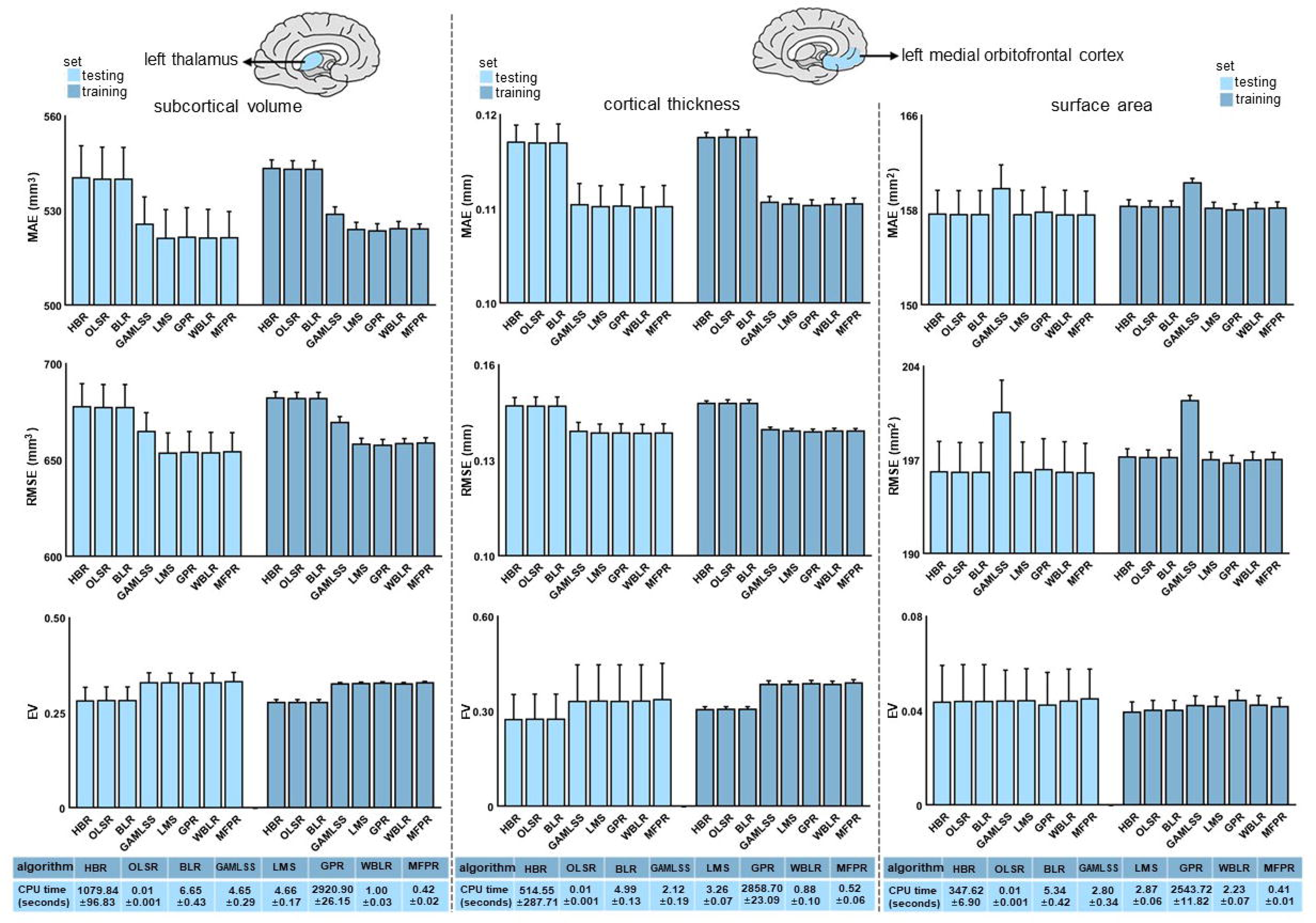
Illustrative examples of comparative algorithm performance. Algorithm performance for each regional morphometric measure was assessed separately in males and females using the mean absolute error (MAE), the root mean square error (RMSE), explained variance (EV), and the central processing unit (CPU) time. The MAE, RMSE, EV, and CPU times of the models for left thalamic volume (left panel), the left medial orbitofrontal cortical thickness (middle panel) and surface area (right panel) as exemplars here for females and in appendix 1, p 4, figure S3 for males. The pattern identified was the same across all region-specific models and in both sexes (appendix 1, pp 5– 6, figures S4–S5). HBR=Hierarchical Bayesian Regression; OLSR=Ordinary Least Squares Regression; BLR=Bayesian Linear Regression; GAMLSS=Generalized Additive Models for Location, Scale and Shape; LMS=Lambda (λ), Mu (μ), Sigma (σ) Quantile Regression; GPR=Gaussian Process Regression; WBLR=Warped Bayesian Linear Regression; MFPR=Multivariate Fractional Polynomial Regression.

*Selection of explanatory variables for model optimization:* We considered the following covariates in all models: scanner vendor, Euler Number (EN), FreeSurfer version, and global neuroimaging measures (i.e., ICV, mean cortical thickness, or mean cortical surface area, as indicated) and their linear and non-linear combinations. In figure 3, we illustrate the effects of the covariates for the MFPR-derived models of the left thalamic volume and left medial orbitofrontal cortical thickness and surface area in females are shown as exemplars (the corresponding data for males in appendix 1, p 7, figure S6). The same pattern was observed for all regions across sex-specific MFPR models as shown in appendix 1 (pp 8– 9, figures S7 and S8) and appendix 4. The impact of the scanner, EN (appendix 1, p 3, figure S2), and FreeSurfer version on model performance was minimal while the opposite was the case for the global neuroimaging measures. Therefore, optimised models included age and global neuroimaging measures (i.e., ICV, mean cortical thickness, or mean cortical surface area, as indicated).

**Figure 3.**
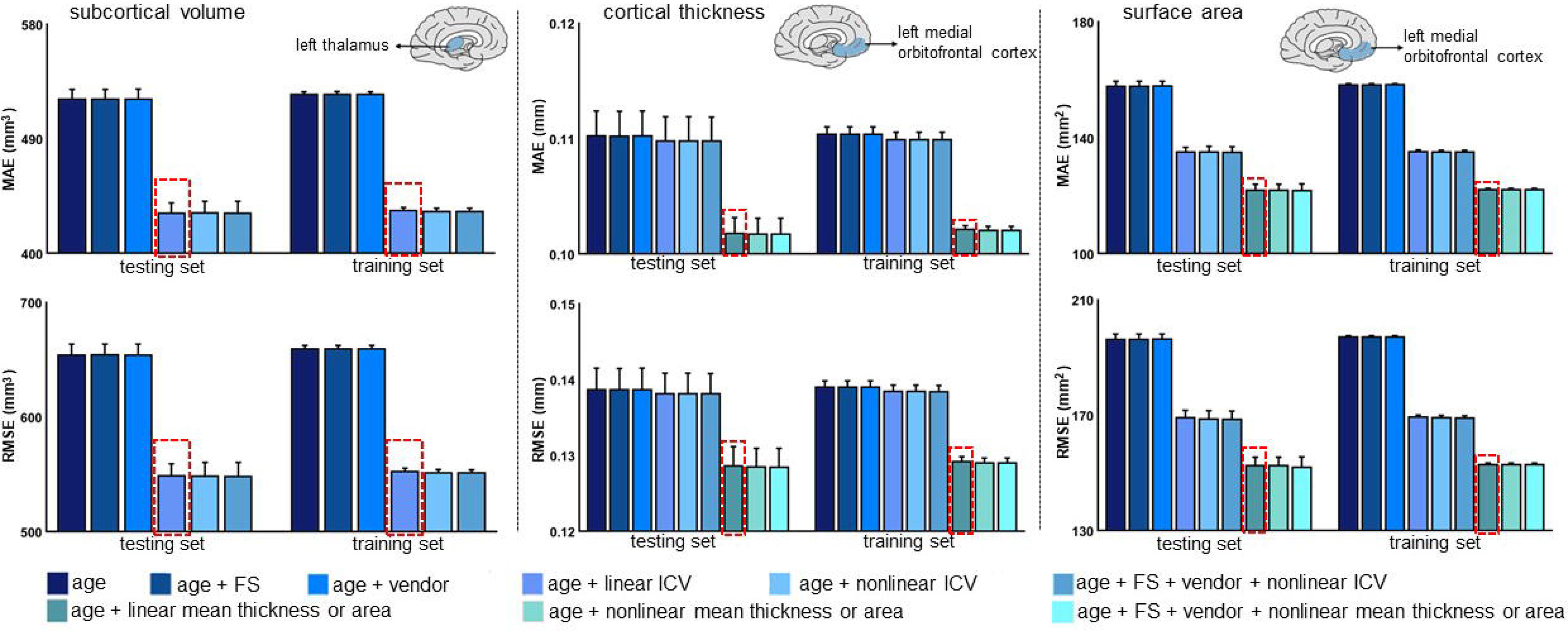
Illustrative examples of the performance of MFPR-derived models as a function of explanatory variables. For each regional morphometric measure, sex-specific models derived from all algorithms were trained and tested using nine different covariate combinations that included effects of age, FreeSurfer version (FS), Euler Number, scanner vendor, intracranial volume (ICV) and global estimates of mean cortical thickness or area. The mean absolute error (MAE) and root mean square error (RMSE) of models for left thalamic volume (left panel), the left medial orbitofrontal cortical thickness (middle panel), and surface area (right panel) derived from Multivariate Fractional Polynomial Regression (MFPR) for females are presented as exemplars; the optimal variable combination is marked with a dashed frame. The corresponding data for males are presented in appendix 1 (p 7, figure S6). The data for other algorithms are shown in appendix 1 (pp 8–12, figures S7–S11 and S2). In both sexes, the pattern identified was identical for all region-specific models.

We then compared the MAE, RMSE, and CPU time for each of the sex- and region-specific optimized models derived from the other algorithms. Statistical comparison of the models from each algorithm at P_FDR_<0·05 indicated comparable performance for the optimised MFPR-, WBLR- and GPR-derived models that outperformed the optimized models derived from the other algorithms. We illustrate these findings for females in figure 4 using the left thalamic volume and left medial orbitofrontal cortical thickness and surface area as exemplars (the corresponding data in males and for all other regions in appendix 1, pp 10–12, figures S9–S11 and appendix 6). In addition to retaining their accuracy, the MFPR-derived models remained the most computationally efficient with CPU times of less than a second. Accordingly, we define as “optimal models” the sex- and region-specific models that were based on the MFPR algorithm with nonlinear fractional polynomials of age and linear effects of the appropriate global neuroimaging measure (i.e., ICV for models of regional subcortical volumes, mean cortical thickness for models of regional cortical thickness and mean cortical surface area for models of regional cortical surface area).

**Figure 4.**
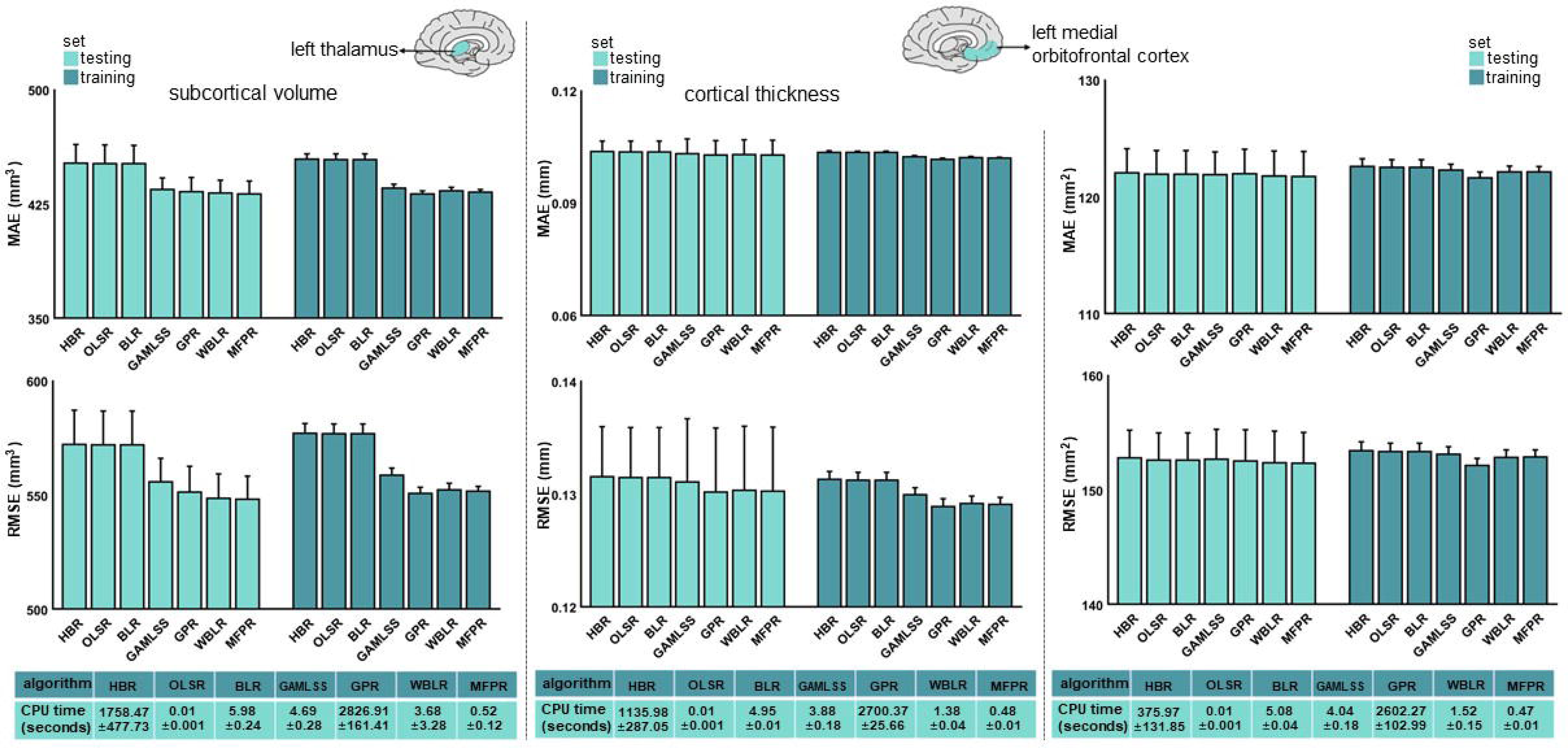
Illustrative examples of the comparative performance of OLSR, BLR, HBR, GPR, GAMLSS, WBLR, and MFPR-derived optimised models. Region-specific models with the optimised covariate combination were estimated in males and females separately using Ordinary Least Squares Regression (OLSR), Bayesian Linear Regression (BLR), Hierarchical Bayesian Regression (HBR), Gaussian Process Regression (GPR), Generalized Additive Models for Location, Scale, and Shape (GAMLSS), Warped Bayesian Linear Regression (WBLR), and Multivariable Fractional Polynomial Regression (MFPR). Model performance was assessed in terms of mean absolute error (MAE), root mean square error (RMSE), and central processing unit (CPU). The MAE, RSME, and CPU time of the models for left thalamic volume (left panel), the left medial orbitofrontal cortical thickness (middle panel), and surface area (right panel) in females are presented as exemplars and in appendix 1 (p 10, figure S9) for males.

*Sensitivity analyses*: (1) Sample size: The MAE and RMSE values of the optimised MFPR-derived sex- and region-specific models plateaued at a sample size of approximately 3,000 participants as shown in figure 5 for females and for males in appendix 1 (p 13, figure S12).

**Figure 5.**
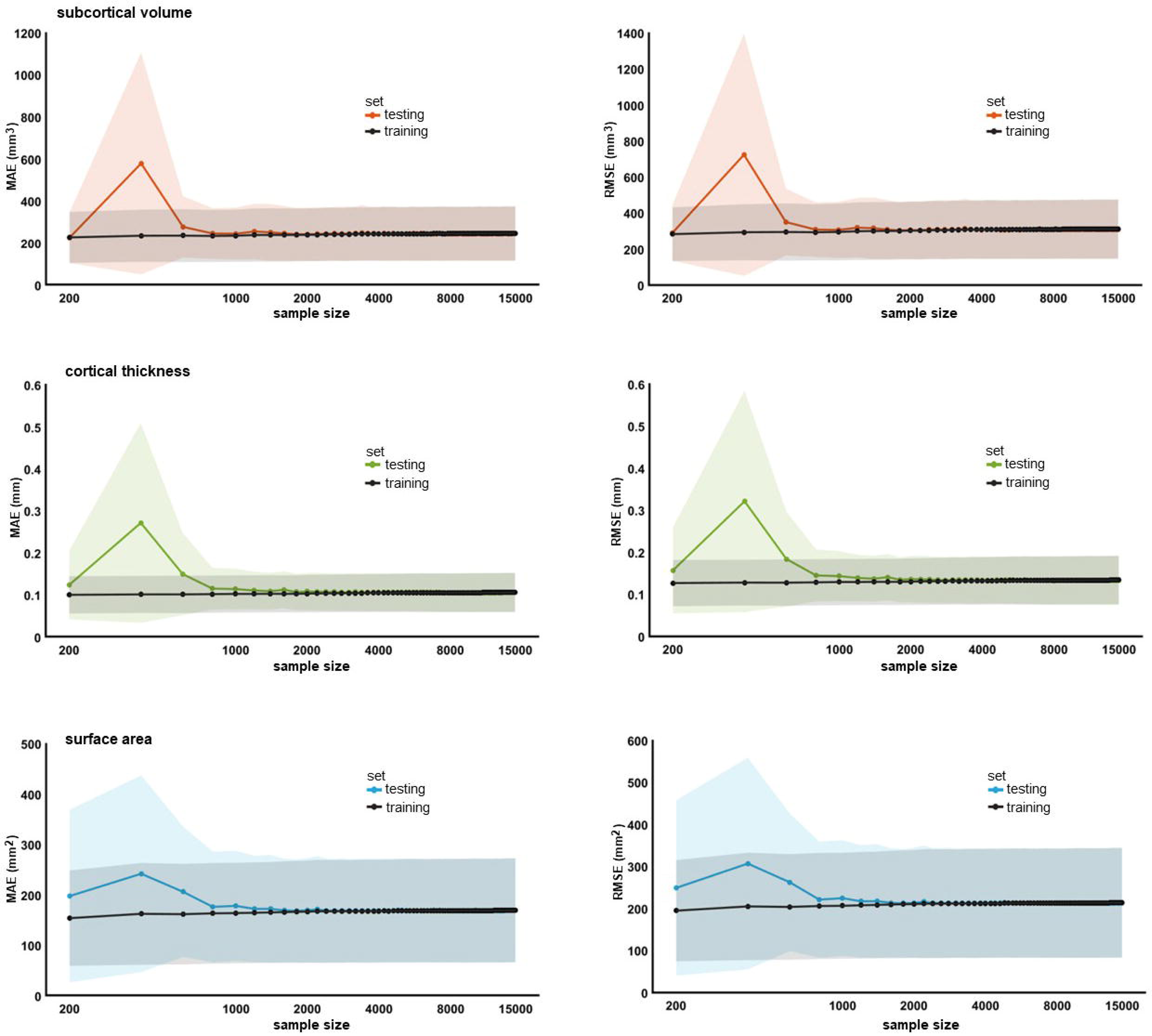
Performance of region-specific MFPR-derived models as a function of sample size. Models for each regional morphometric measure were estimated in random sex-specific subsets of 200 to 15,000 participants, in increments of 200, generated from the study sample. Each line represents the values of the mean absolute error (MAE), or root mean square error (RMSE) derived from the optimized Multivariate Fractional Polynomial Regression (MFPR) models of all regional morphometric measure as a function of sample size; shadowed area represents the standard deviation. The pattern identified was identical in both sexes. The data for females are shown here and for males in appendix 1 (p 13, figure S12).

(2) Age bins: The MAE and RMSE values of the optimised MFPR-derived sex- and region-specific models in each of the nine age bins are presented in figure 6 for females and for males in appendix 1 (p 14, figure S13) and appendix 5. Across all age bins, the correlation coefficient between the MAE or RMSE values of the sex- and region-specific models obtained from the full study sample and MAE or RMSE values of the corresponding models estimated in each age bin were all greater than 0·98, suggesting the robustness of the model accuracy across all age groups.

**Figure 6.**
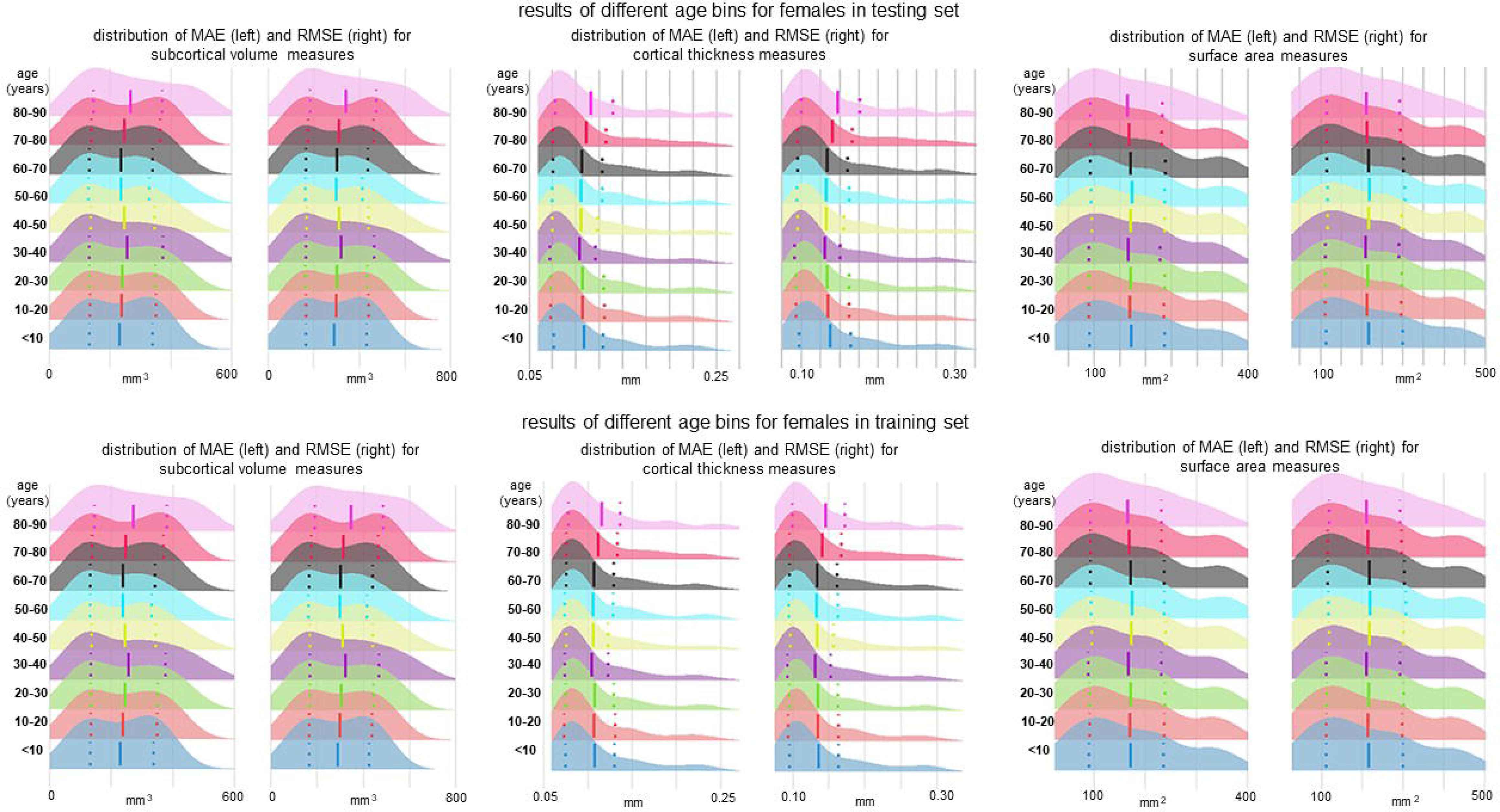
Performance of region-specific models in distinct age bins. Sex- and region-specific models of all morphometric measures for different age bins were estimated by partitioning the sex-specific training and testing subsets of the study sample into nine age bins (i.e., age≤10 years; 10<age≤20 years; 20<age≤30 years; 30<age≤40 years; 40<age≤50 years; 50<age≤60 years; 60<age≤70 years; 70<age≤80 years; 80<age≤90 years). Details are provided in appendix 5. The figure presents the distribution of the mean absolute error (MAE) and the root mean square error (RMSE) across all region-specific models in females in the training (upper panel) and testing subset (lower panel). The pattern was identical in both sexes and the results for males are presented in appendix 1 (p 14, figure S13).

(3) GAMLSS: Comparison of different GAMLSS models and software confirmed the superiority of the choice reported here compared to other alternatives (appendix 1, pp 15–19 figures S14–S17).

(4) Effect of site: The performance of the OLSR, BLR, GAMLSS, WBLR, HBR, and MFPR was compared when site was modeled either as a random factor or by harmonisation using ComBat-GAM. These comparisons excluded the LMS as it does not accommodate multiple explanatory variables and the GPR because it assumes only continuous variables. The top-performing algorithm when the site was used as a random effect was still the MFPR, followed closely by WBLR. Further, the model performance of the MFPR algorithm in terms of MAE was similar regardless of how site was handled. Details in appendix 1, p 20, figure S18.

(5) Optimised MFPR model performance in longitudinal datasets: In figure 7, we demonstrate the stability of the regional Z-scores derived with the optimised MFPR models applied to structural MRI data of healthy participants in the SLIM and QTAB samples scanned with an average interval of approximately two years.

**Figure 7.**
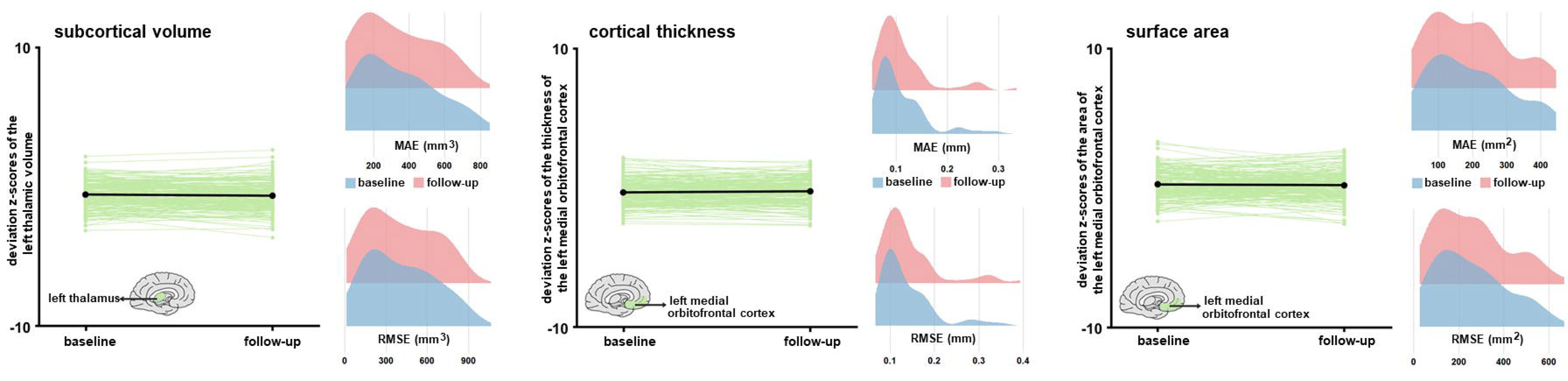
Stability of the normative deviation scores (Z-scores) in longitudinal neuroimaging data. We illustrate the stability of the optimized MFPR-derived models over an average interval of two years in data from the SLIM and QTAB samples using the left thalamic volume (left panel), the left medial orbitofrontal cortical thickness (middle panel), and surface area (right panel) as exemplars. Within each panel, the left-hand figure shows the Z-scores of each participant at baseline and follow-up and the right-hand figure shows the distribution of the mean absolute error (MAE) and root mean squared error (RMSE) at baseline and follow-up. SLIM=Southwest Longitudinal Imaging Multimodal Study; QTAB= Queensland Twin Adolescent Brain Study.

### Relevance of normative models of brain morphometry for mental illness

In the HCP-EP cohort, the accuracy of the diagnostic classification of the SVCs that used regional Z-scores performed similarly regardless of the normative model and outperformed the SVC with the observed data. In figure 8, we illustrate these findings, by showing that the SVC which used Z-scores from the optimized MFPR models achieved an AUC of 0·63 (p<0·001) while the accuracy of the SVC that used observed data was indistinguishable from chance (AUC 0·49). Information on other models in appendix 1, p 21, figure S19.

**Figure 8.**
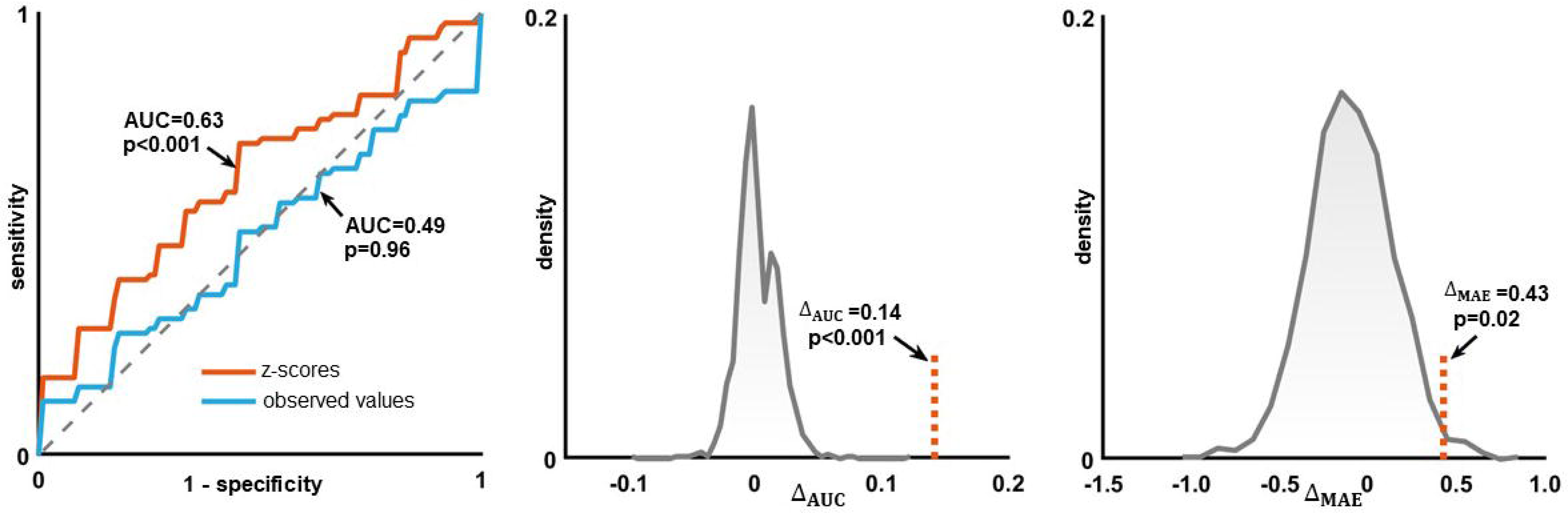
Accuracy of diagnostic classification and accuracy of psychotic symptom prediction using brain regional normative deviation scores or observed neuromorphometric data. Left panel: Diagnostic classification accuracy in the Human Connectome Project -Early Psychosis (HCP-EP) sample. Receiver Operating Characteristic (ROC) curves of the models distinguishing patients from controls using the observed regional neuromorphometric measures (blue curve) or the deviation Z-scores from the normative model (orange curve); Middle panel: the area under the curve (AUC) difference between a support vector machine classifier using the observed regional neuromorphometric measures and another using regional normative deviation scores (Z-scores) derived from the optimised multivariate fractional polynomial regression (MFPR) model was examined through 1000 permutations. The AUC difference is marked by a vertical dotted line. Right panel: Predictive Accuracy of Psychotic Symptoms in the HCP-EP sample. The mean absolute error (MAE) difference between a ridge regression using the observed regional neuromorphometric measures and another using Z-scores derived from the optimised MFPR model was examined through 1000 permutations. The MAE difference is marked by a vertical dotted line. Information on other models in appendix 1, p 22, figure S19.

The predictive accuracy for psychotic symptom severity of the ridge regression models using the Z-scores from the different normative models performed similarly to each other and to the model using observed data; none achieved above chance performance (appendix 1, p 21, figure S19). In figure 8, we illustrate these findings by showing the predictive accuracy of the regression models using optimised MFPR-derived Z-scores or observed data.

## Discussion

The present study undertook a comparative evaluation of eight algorithms commonly used for normative modeling using morphometric data from a multisite sample of 37,407 healthy individuals. Sex-specific models based on the MFPR algorithm with nonlinear fractional polynomials of age and linear global neuroimaging measures emerged as optimal based on their performance and computational efficiency, the latter being an important consideration when analysing large datasets. These models were robust to variations in sample composition with respect to age and their performance plateaued at sample sizes of ≈3,000. The optimised sex-specific MFPR models showed longitudinal stability over an average interval of two years and the Z-scores derived from these models outperformed observed neuromophometric measures in distinguishing patients with psychosis from healthy individuals.

The presented findings validate our choice to use MFPR in our previous normative studies on brain morphometry^2,3^ and white matter microstructure based on diffusion-weighted MRI.^25^ Further, after testing the impact of multiple combinations of explanatory variables on model performance we found that global morphometric measures (i.e., ICV, mean cortical thickness, mean cortical surface area) had the most significant effect. This observation is aligned with prior literature on the contribution of ICV in explaining the variance of regional subcortical volumes and cortical surface area measures.^26, 27^ The present study extended these findings by demonstrating that mean cortical thickness and mean surface area outperformed ICV as explanatory variables in normative models of regional cortical thickness and cortical surface area respectively. Accordingly, the optimal normative models for brain morphometry consisted of an MFPR algorithm and a combination of explanatory variables that comprised nonlinear fractional polynomials of age and linear global measures of ICV, mean cortical thickness, and mean cortical surface area for models of regional subcortical volume, regional cortical thickness, and regional cortical surface area respectively. Sensitivity analyses across different age bins supported the applicability of the models developed in the whole study sample, which spanned an age range of 3–90 years, to groups with a more restricted age range and at different points in their life trajectories. The optimised sex-specific MFPR models showed longitudinal stability over an average follow-up period of two years as would be expected for healthy adults over relatively short time periods.^1–3^

Site variation is a major challenge when aggregating multisite data as it may confound or bias results. The most common methods for minimizing site effects involve either site harmonization using ComBat-GAM prior to normative modeling or the inclusion of site as an explanatory variable in the normative models. A recent publication that used a smaller sample (569 healthy participants) and a narrower age range (6–40 years) suggested the HBR with site as an explanatory variable may be superior to ComBat-based site harmonization for the normative modeling of brain morphometry.^12^ We found no support for this assertion in our sensitivity analyses. An additional advantage of using ComBat-GAM is that it removes the requirement for calibration and model parameter adaptation every time the model is applied to data from a new site. In the HBR models, by contrast, pre-trained parameters can be used for new data if they originate from one of the sites in the training dataset^10^ or under the assumption that the variation accounted for by an unseen site should align with that of the sites in the training dataset.^12^

Prior studies have shown that sex accounts for a significant amount of variance in brain morphology, both cross-sectionally^16^ and longitudinally.^28^ Accordingly, we developed sex-specific models for each brain morphometric measure thus extending prior normative studies that considered males and females together.^1,13^ Additionally, we provide normative models for regional cortical surface area measures that were not included in prior studies^1,13^ despite the important functional implications of age-related changes in the cortical surface area for cognition during development and aging. We note that the current normative model is compiled cross-sectionally, from people of different ages who experienced different exposures to factors that can influence brain health. In later life, samples of healthy individuals are likely to include those that are more resilient to mortality and morbidity.

There are several methodological limitations pertinent to the present study. Specifically, our study could benefit from the inclusion of more young and middle-aged adults and data from longitudinal follow-up over long periods of time. Testing the generalizability of our models to populations with specific ancestries is an important next step. We did not include an exhaustive list of potential explanatory variables. It could be argued that the inclusion of other variables, such as childhood adversity, premature birth, or socioeconomic status that are known to influence brain morphometry,^29,30^ could have further improved model performance. Exploring this possibility further could be best achieved within the context of single large-scale studies where such variables would be consistently recorded in all participants. On the other hand, the inclusion of multiple explanatory variables in the normative model itself could restrict its applicability to those datasets where all such features were assessed.

In conclusion, this study presents a detailed evaluation of the comparative performance of the key eight algorithms used for normative modeling and of the influence of key parameters pertaining to site effects, covariates, sample size, and sample composition with respect to age on model accuracy and robustness. Based on the evidence provided, we consider the sex-specific optimised MFPR models developed here to be advantageous in terms of accuracy and efficiency compared to other options. We therefore provide these models in a user-friendly web platform (www.centilebrain.org) that enables the estimation of normative deviation scores from any sample with minimal technical and computing requirements.

## Data availability

Access to individual participant data from each dataset is available through access requests addressed to the principal investigators of the original studies or to the relevant data repositories. Details are provided in appendix 2.

## Code availability

A dedicated web portal (https://centilebrain.org) provides the optimal model parameters, as pre-trained models, to be applied to any user-specified dataset in the context of open science.

## Author contributions

All authors contributed to data collection, data interpretation, and manuscript writing. In addition, authors RG, YY, YXQ, YF, SC, CG, SSH, FN, and SF were directly involved with data analysis and the comparison of the normative models.

## Acknowledgments

**Funding:** EU Seventh Framework Programme: 278948, 602450, 603016, 602805; 602450; EU Horizon 2020 Programme: 667302, 643051; European Research Council: ERC- 230374; EU Joint Programme-Neurodegenerative Disease Research: FKZ:01ED1615;

Australia: Australian National Health and Medical Research Council: 496682, 1009064;

Germany: Federal Ministry of Education and Research: 01ZZ9603, 01ZZ0103, 01ZZ0403;

Netherlands: Vici Innovation Program:91619115, 016- 130- 669; Nederlandse Organisatie voor Wetenschappelijk Onderzoek (NWO): Cognition Excellence Program: 433- 09- 229, NW0-SP 56- 464- 14192, NWO- MagW 480- 04- 004, NWO 433- 09- 220, NWO 51-02-062, NWO 51-02-061; Organization for Health Research and Development: 480- 15- 001/674, 024-001-003, 911- 09- 032, 056- 32- 010, 481- 08- 011, 016- 115- 035, 31160008, 400- 07- 080, 400- 05- 717, 451- 04- 034, 463- 06- 001, 480- 04- 004, 904- 61- 193, 912- 10- 020, 985- 10- 002, 904- 61- 090, 912- 10- 020, 451- 04- 034, 481- 08- 011,, 056- 32- 010, 911- 09- 032; Dutch Health Research Council: 10- 000- 1001; Biobanking and Biomolecular Resources Research Infrastructure: 184-033-111, 84.021.00;

Norway: Research Council of Norway: 223273; South and Eastern Norway Regional Health Authority: 2017- 112, 2019-107, 2014- 097, 2013- 054;

Russian Federation: Russian Foundation for Basic Research: 20- 013- 00748;

Spain: Fundación Instituto de Investigación Marqués de Valdecilla: API07/011, NCT02534363, NCT0235832; Instituto de Salud Carlos III: PI14/00918, PI14/00639, PI060507, PI050427, PI020499;

Sweden: Swedish Research Council: 523- 2014- 3467, 2017- 00949, 521- 2014- 3487, K2007- 62X- 15077- 04- 1, K2008- 62P- 20597- 01- 3, K2010- 62X- 15078- 07- 2, K2012- 61X-15078- 09- 3; Knut and Alice Wallenberg Foundation;

UK: Medical Research Council: G0500092;

USA: National Institutes of Health/Mental Health/Aging/ Child Health and Human Development/ Drug Abuse/National Center for Advancing Translational Sciences: UL1 TR000153; U24RR025736- 01, U24RR021992, U54EB020403, U24RR025736, U24RR025761, P30AG10133, R01AG19771, R01MH117014, R01MH042191; R01HD050735, 1009064, 496682; R01MH104284, R01MH113619, R01MH116147 R01MH116147, R01MH113619, R01MH104284, R01MH090553, R01MH090553, R01CA101318; RC2DA029475, T32MH122394

We thank Dr. Andre F. Marquand and Dr. Seyed Mostafa Kia, both from Radboud University, the Netherlands, for their guidance with the HBR models. This work was supported by the computational resources and staff expertise provided by the Advanced Research Computing at the University of British Columbia and by the Scientific Computing at the Icahn School of Medicine at Mount Sinai (supported by the Clinical and Translational Science Awards grant UL1TR004419 from the National Center for Advancing Translational Sciences).

