## Appendix 1 for "Normative Modeling of Brain Morphometry Across the Lifespan Using CentileBrain: Algorithm Benchmarking and Model Optimization"

**Table of Contents**

**Appendix 1**
**S1. Samples** **2**
Supplementary Figure S1. Age-by-sex distribution of the study sample 2
**S2. Quality assessment of neuroimaging data** **3**
Supplementary Figure S2. Illustrative results of the minimal effect of Euler Number on the performance of the Multivariate Fractional Polynomial Regression models. 3
**S3. Comparative evaluation of algorithms** **4**
Supplementary Figure S3. Illustrative example of comparative algorithm performance for males 4
Supplementary Figure S4. Comparative algorithm performance of region-specific models in females 5
Supplementary Figure S5. Comparative algorithm performance of region-specific models in males 6
**S4. Selection of explanatory variables for model optimization** **7**
Supplementary Figure S6. Illustrative examples of the performance of MFPR-derived models as a function of explanatory variables for males 7
Supplementary Figure S7. Performance of region-specific MFPR-derived models as a function of explanatory variables in females 8
Supplementary Figure S8. Performance of region-specific MFPR-derived models as a function of explanatory variables in males 9
Supplementary Figure S9. Illustrative examples of the comparative performance of HBR, OLSR, BLR, GAMLSS, GPR, WBLR, and MFPR-derived models in males 10
Supplementary Figure S10. Comparative performance of all region-specific optimized HBR, OLSR, BLR, GAMLSS, GPR, WBLR, and MFPR-derived models in females 11
Supplementary Figure S11. Comparative performance of all region-specific optimized HBR, OLSR, BLR, GAMLSS, GPR, WBLR, and MFPR-derived models in males 12
**S5. Sensitivity analyses** **13**
**S5.1 Sample size** **13**
Supplementary Figure S12. Performance of region-specific MFPR-derived models as a function of sample size for males 13
**S5.2 Age bins** **14**
Supplementary Figure S13. Performance of region-specific models in distinct age groups for males 14
**S5.3 Sensitivity analyses pertaining to the GAMLSS** **16**
Supplementary Figure S14. Comparative evaluations of five GAMLSS models in females 17
Supplementary Figure S15. Comparative evaluations of five GAMLSS models in males 18
Supplementary Figure S16. Comparative evaluations of GAMLSS models using the "caret" and "gamlss" software in females 19
Supplementary Figure S17. Comparative evaluations of GAMLSS models using the "caret" and "gamlss" software in males 20
**S5.4 Sensitivity analyses pertaining to site** **21**
Supplementary Figure S18. Effect of site handling on algorithm performance 21
**S6. Clinical relevance of normative brain models for mental illness** **22**
Supplementary Figure S19. Comparative accuracy of diagnostic classification and symptom severity prediction in psychosis using either brain regional Z-scores or observed neuromorphometric data of the Human Connectome Project-Early Psychosis cohort 22
**S7. List of appendices 2-6 in Excel format** **23**
**Supplementary References** **24**

**S1. Samples**
We collated data from 87 datasets. The age-by-sex distribution of the current study sample is shown in supplementary figure S1 and details of the contributing datasets are provided in appendix 2. In the Figure below, we also illustrate the geographic distribution of the sample, with different colors representing the geographical region of each site. The clustering was conducted on all 150 morphometric measures (68 cortical thickness measure, 68 surface area measures, and 14 subcortical volume measures) of the entire sample. We did not identify region-specific clusters. The Figure was generated using t-SNE (t-distributed stochastic neighbor embedding) plot to visualize the high dimensional data into a 2D plot.

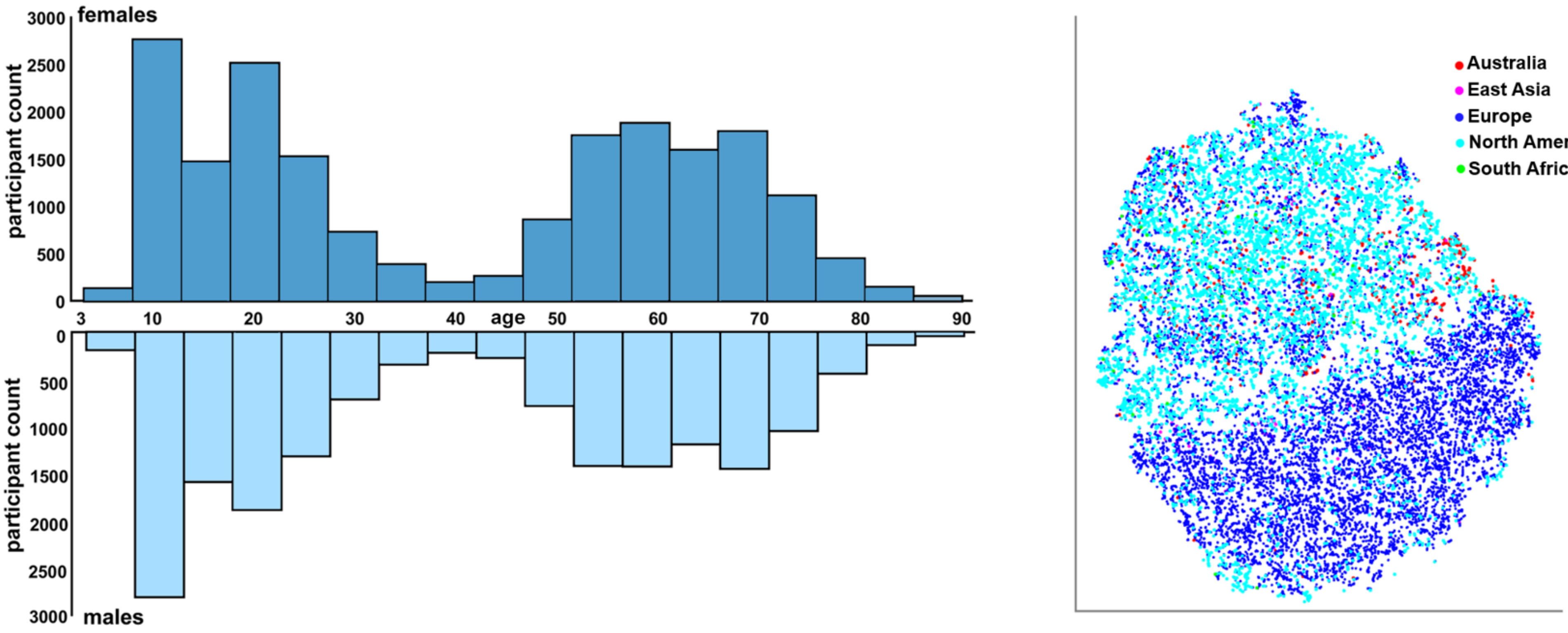


**Supplementary Figure S1. (left panel) Age-by-sex distribution of the study sample, and (right panel) geographic distribution of the sample.**

**S2. Quality assessment of neuroimaging data**

For samples participating in the ENIGMA Lifespan Working Group (appendix 2), quality assessment followed the ENIGMA consortium pipeline (http://enigma.ini.usc.edu/protocols/imaging-protocols/).[1–7] These pipelines were therefore applied to 18,787 individuals in the study sample. Data from the Adolescent Brain Cognitive Development study were considered high-quality based on their quality assessment protocols (Txt file= FreeSurfer QC; item= fsqc_qc; scan excluded if 0).[8] This applied to 3,116 individuals in the study sample. For the remainder of the sample involving 15,504 participants, the T1-weighted images were downloaded and segmented locally, and the results were quality assessed using Qoala-T tool.[9]

As an additional assessment of image quality, we calculated the Euler number (EN), a reliable indicator that closely approximates the manual assessment of scan quality[10, 11] for the entire dataset. The addition of the EN to the covariates did not make a measurable difference in any model. We illustrate that for the Multivariate Fractional Polynomial Regression (MFPR) models for the left thalamic volume in males using the UKB data which are publicly accessible. The pattern observed for all other regions and models in each sex is the same and the data are available upon request.

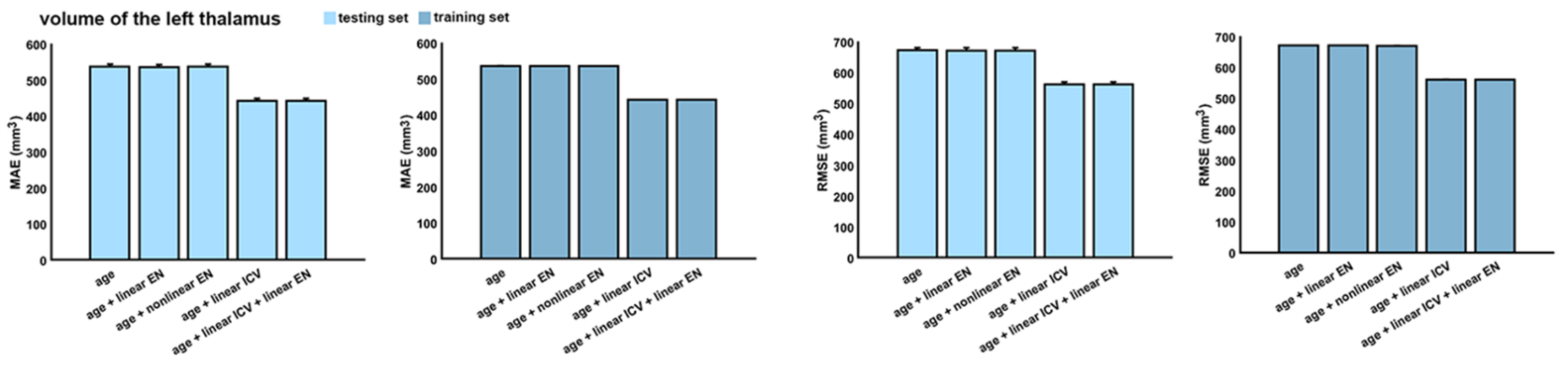


**Supplementary Figure S2. Illustrative results of the minimal effect of Euler Number on the performance (MAE; mean absolute error and RMSE; root mean square error) of the Multivariate Fractional Polynomial Regression models.**

**S3. Comparative evaluation of algorithms**

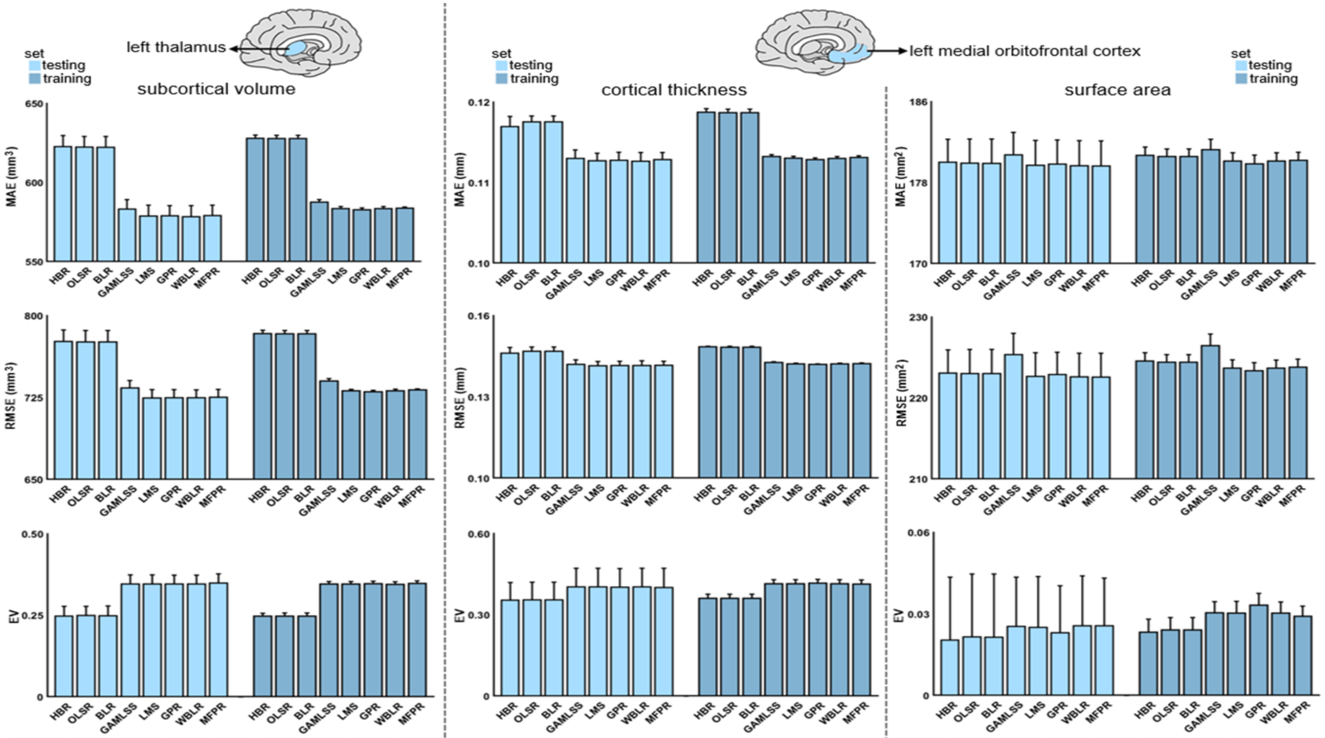


| algorithm | HBR | OLSR | BLR | GAMLSS | LMS | GPR | WBLR | MFPR |
|---|---|---|---|---|---|---|---|---|
| CPU time (seconds) | 651.29 ±140.58 | 0.01 ±0.01 | 5.82 ±0.37 | 3.16 ±0.06 | 3.85 ±0.10 | 1814.16 ±8.27 | 0.88 ±0.07 | 0.42 ±0.04 |

| algorithm | HBR | OLSR | BLR | GAMLSS | LMS | GPR | WBLR | MFPR |
|---|---|---|---|---|---|---|---|---|
| CPU time (seconds) | 493.18 ±236.18 | 0.01 ±0.001 | 4.43 ±0.22 | 1.92 ±0.24 | 2.71 ±0.02 | 1651.53 ±45.08 | 0.82 ±0.04 | 0.34 ±0.03 |

| algorithm | HBR | OLSR | BLR | GAMLSS | LMS | GPR | WBLR | MFPR |
|---|---|---|---|---|---|---|---|---|
| CPU time (seconds) | 128.81 ±14.02 | 0.06 ±0.13 | 5.02 ±0.98 | 3.69 ±0.29 | 2.76 ±0.41 | 1779.09 ±81.07 | 0.79 ±0.04 | 0.40 ±0.07 |

**Supplementary Figure S3. Illustrative example of comparative algorithm performance for males.** Algorithm performance for each regional morphometric measure was assessed separately in males and females using the mean absolute error (MAE), the root mean square error (RMSE), the Explained Variance (EV), and the central processing unit (CPU) time. The MAE, RMSE, EV, and CPU times of the models for left thalamic volume (left panel), the left medial orbitofrontal cortical thickness (middle panel), and surface area (right panel) as exemplars are presented here for males and in figure 2 in the main text for females. HBR=Hierarchical Bayesian Regression; OLSR=Ordinary Least Squares Regression; BLR=Bayesian Linear Regression; GAMLSS=Generalized Additive Models for Location, Scale, and Shape; LMS=Lambda (λ), Mu (μ), Sigma (σ) Quantile Regression; GPR=Gaussian Process Regression; WBLR=Warped Bayesian Linear Regression; MFPR=Multivariable Fractional Polynomial Regression.

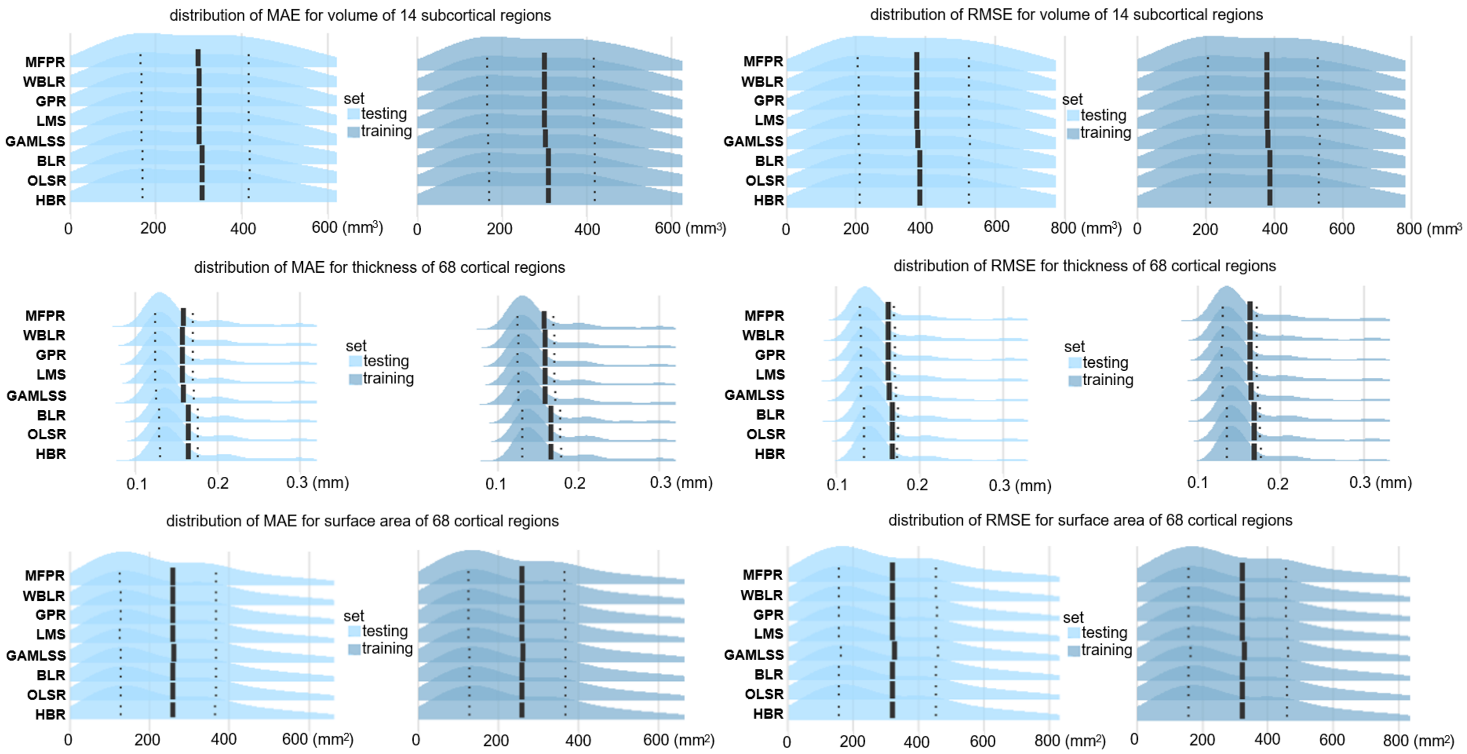


**Supplementary Figure S4. Comparative algorithm performance of region-specific models in females.** The figure presents the distribution of the mean absolute error (MAE) and the root mean square error (RMSE) of all region-specific models. Mean MAE and RMSE values are marked with solid vertical lines, and the 25th percentiles and 75th percentiles are marked with dotted vertical lines. HBR=Hierarchical Bayesian Regression; OLSR=Ordinary Least Squares Regression; BLR=Bayesian Linear Regression; GAMLSS=Generalized Additive Models for Location, Scale, and Shape; LMS=Lambda (λ), Mu (μ), Sigma (σ) Quantile Regression; GPR=Gaussian Process Regression; WBLR=Warped Bayesian Linear Regression; MFPR=Multivariable Fractional Polynomial Regression.

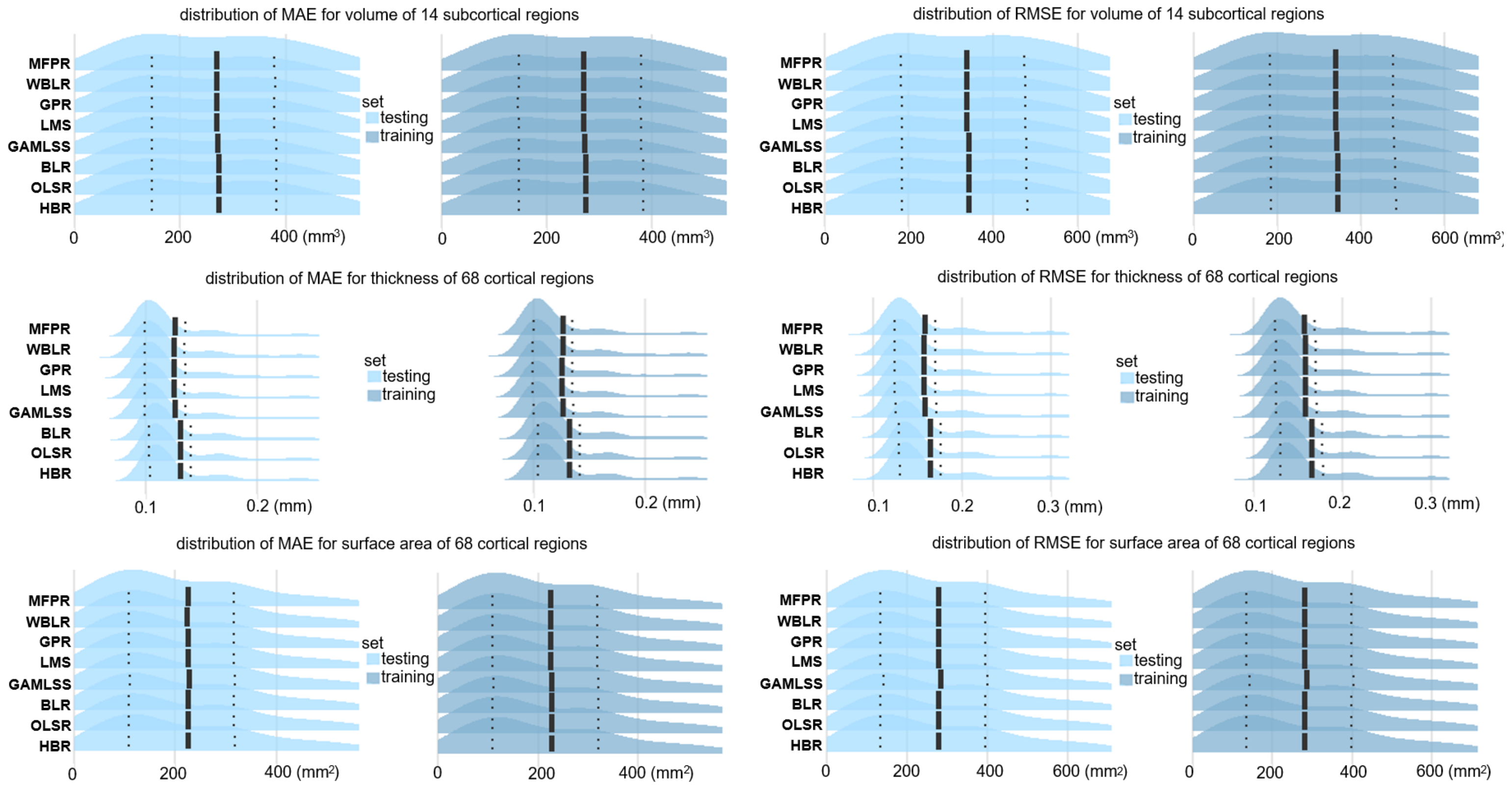


**Supplementary Figure S5. Comparative algorithm performance of region-specific models in males.** The figure presents the distribution of the mean absolute error (MAE) and the root mean square error (RMSE) of all region-specific models. Mean MAE and RMSE values are marked with solid vertical lines, and the 25th percentiles and 75th percentiles are marked with dotted vertical lines. HBR=Hierarchical Bayesian Regression; OLSR=Ordinary Least Squares Regression; BLR=Bayesian Linear Regression; GAMLSS=Generalized Additive Models for Location, Scale, and Shape; LMS=Lambda (λ), Mu (μ), Sigma (σ) Quantile Regression; GPR=Gaussian Process Regression; WBLR=Warped Bayesian Linear Regression; MFPR=Multivariable Fractional Polynomial Regression.

### S4. Selection of explanatory variables for model optimization

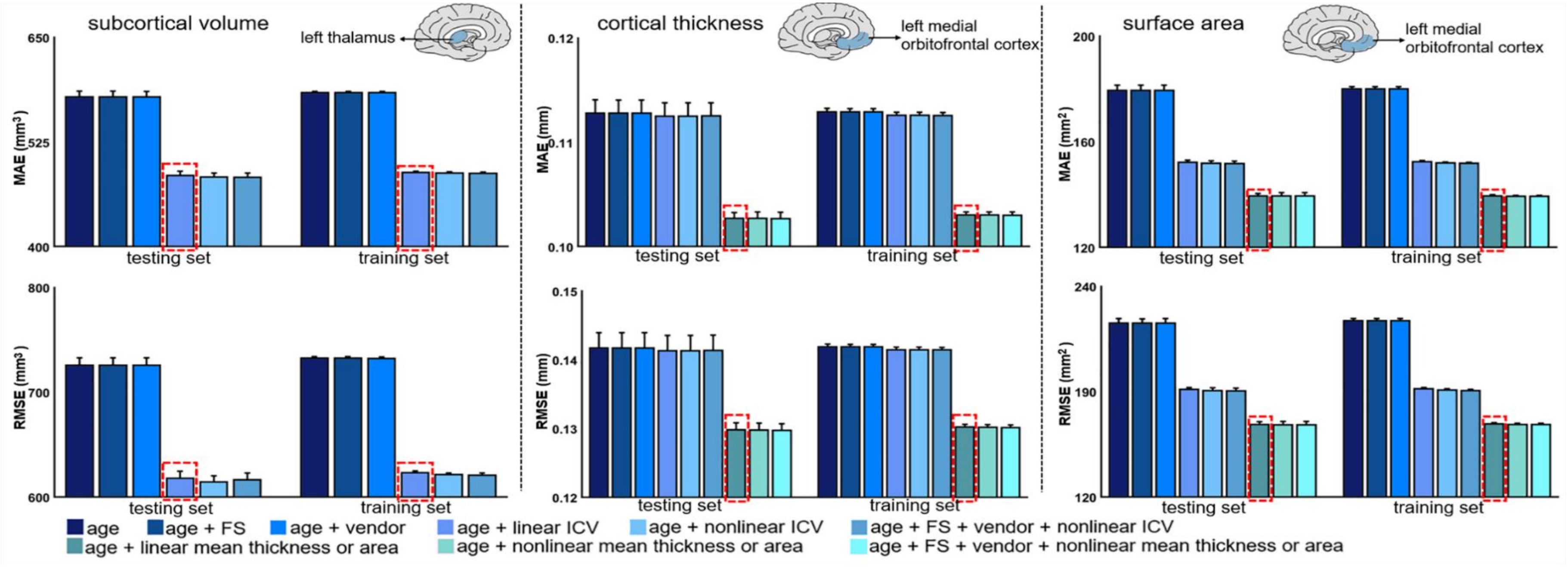


**Supplementary Figure S6. Illustrative examples of the performance of MFPR-derived models as a function of explanatory variables for males.** For each regional morphometric measure, sex-specific models derived from Multivariable Fractional Polynomial Regression (MFPR) were trained and tested using nine different covariate combinations that included linear and non-linear effects of age, FreeSurfer version (FS), scanner vendor, intracranial volume (ICV), and global estimates of mean cortical thickness or area. The mean absolute error (MAE) and root mean square error (RMSE) of all the models in males and females are shown in supplementary figures S7–S8 and appendix 4. In both sexes, the pattern identified was identical for all region-specific models. The MAE and RMSE of the models for left thalamic volume (left panel), the left medial orbitofrontal cortical thickness (middle panel) and surface area (right panel) as exemplars are presented here for males and in figure 3 in the main text for females. The optimal variable combination is marked with a dashed frame.

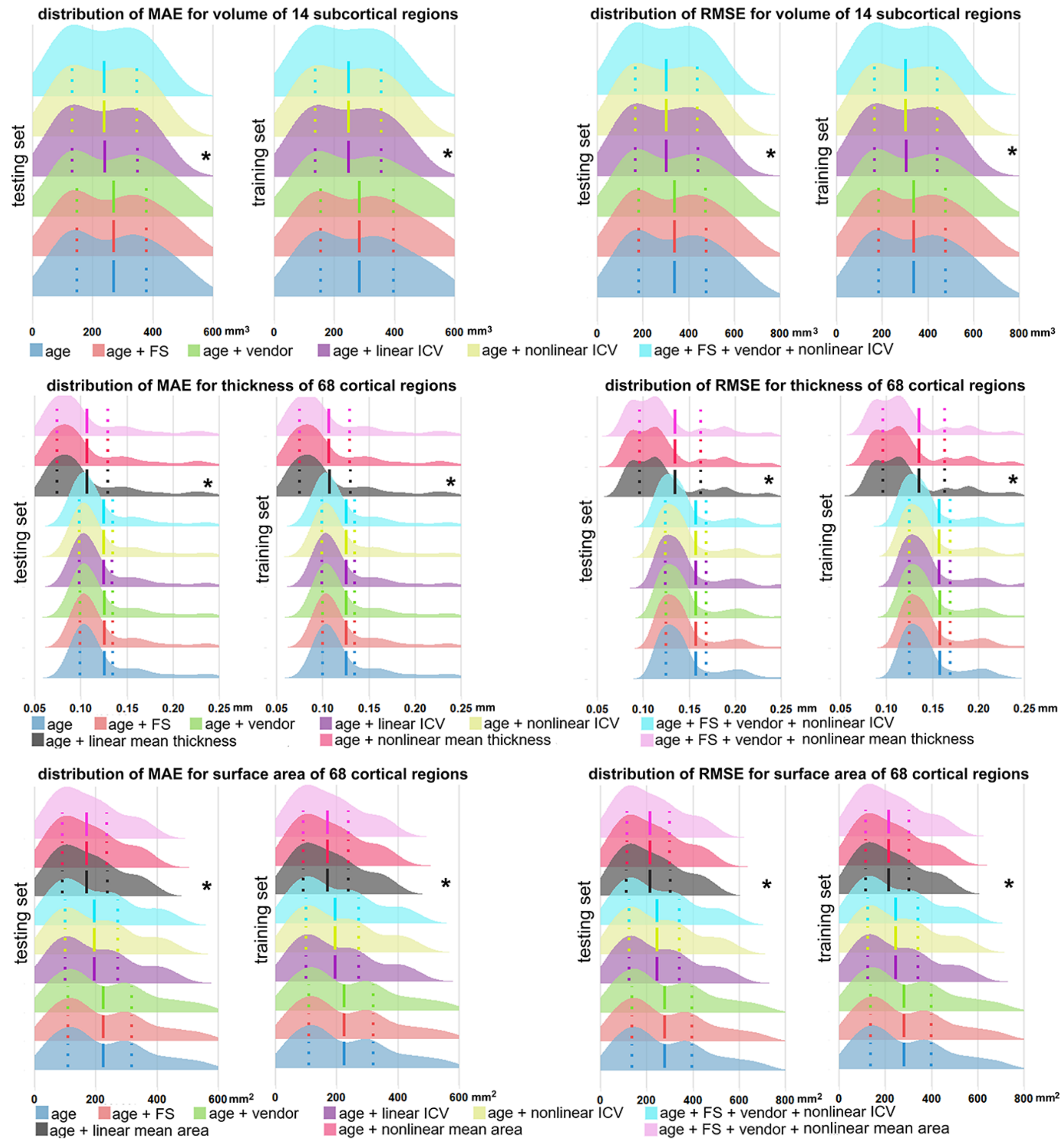


**Supplementary Figure S7. Performance of region-specific MFPR-derived models as a function of explanatory variables in females.** The figure presents the distribution of the mean absolute error (MAE) and the root mean square error (RMSE) for all region-specific models derived from Multivariable Fractional Polynomial Regression (MFPR) as a function of explanatory variables in females. The mean MAE and RMSE values are marked with solid vertical lines, and the 25th percentiles and 75th percentiles are marked with dotted vertical lines. The optimal variable combination is marked with an asterisk. FS=FreeSurfer version; ICV=intracranial volume.

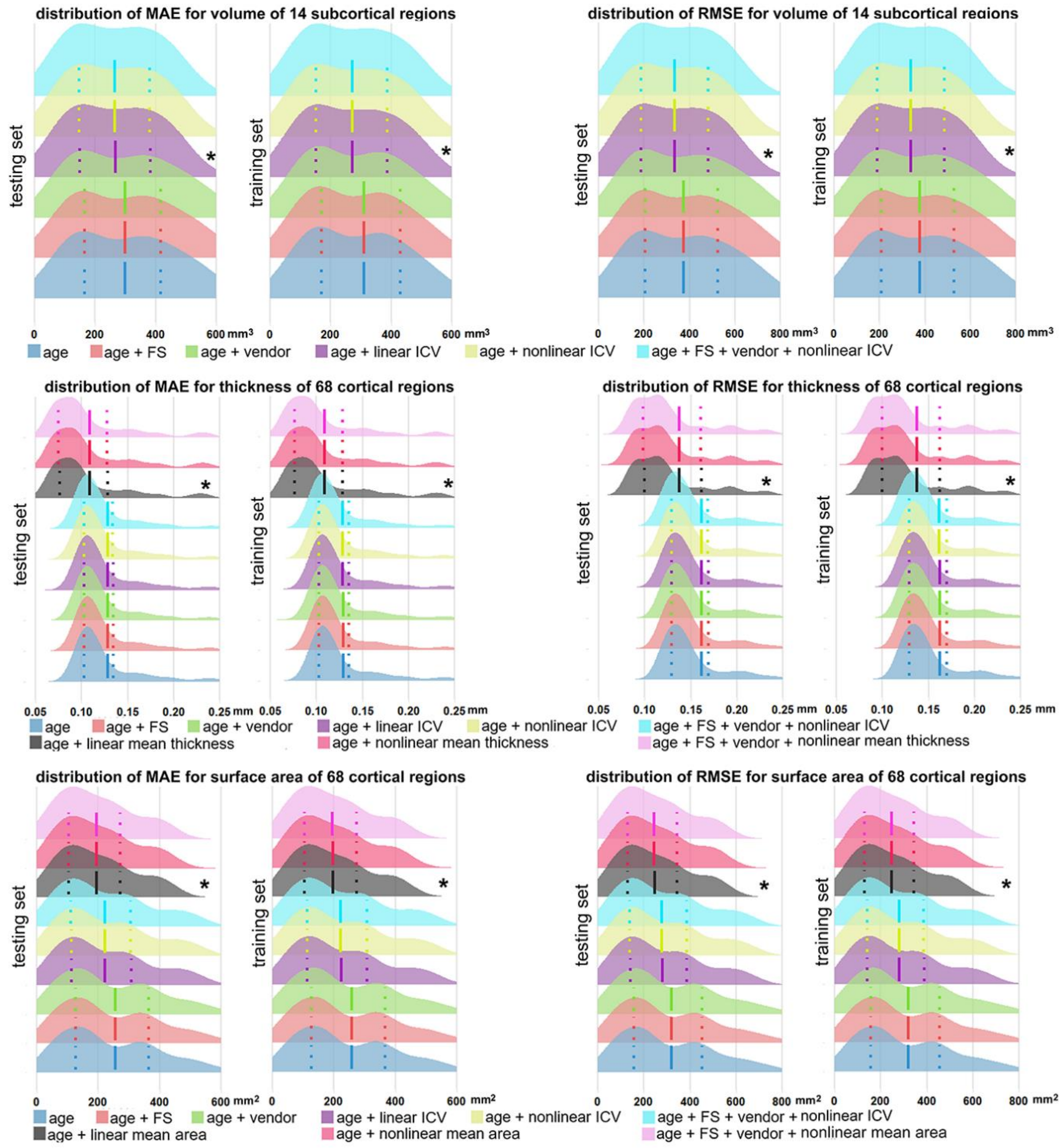


**Supplementary Figure S8. Performance of region-specific MFPR-derived models as a function of explanatory variables in males.** The figure presents the distribution of the mean absolute error (MAE) and the root mean square error (RMSE) for all region-specific models derived from Multivariable Fractional Polynomial Regression (MFPR) as a function of explanatory variables in males. The mean MAE and RMSE values are marked with solid vertical lines, and the 25th percentiles and 75th percentiles are marked with dotted vertical lines. The optimal variable combination is marked with an asterisk. FS=FreeSurfer version; ICV=intracranial volume.

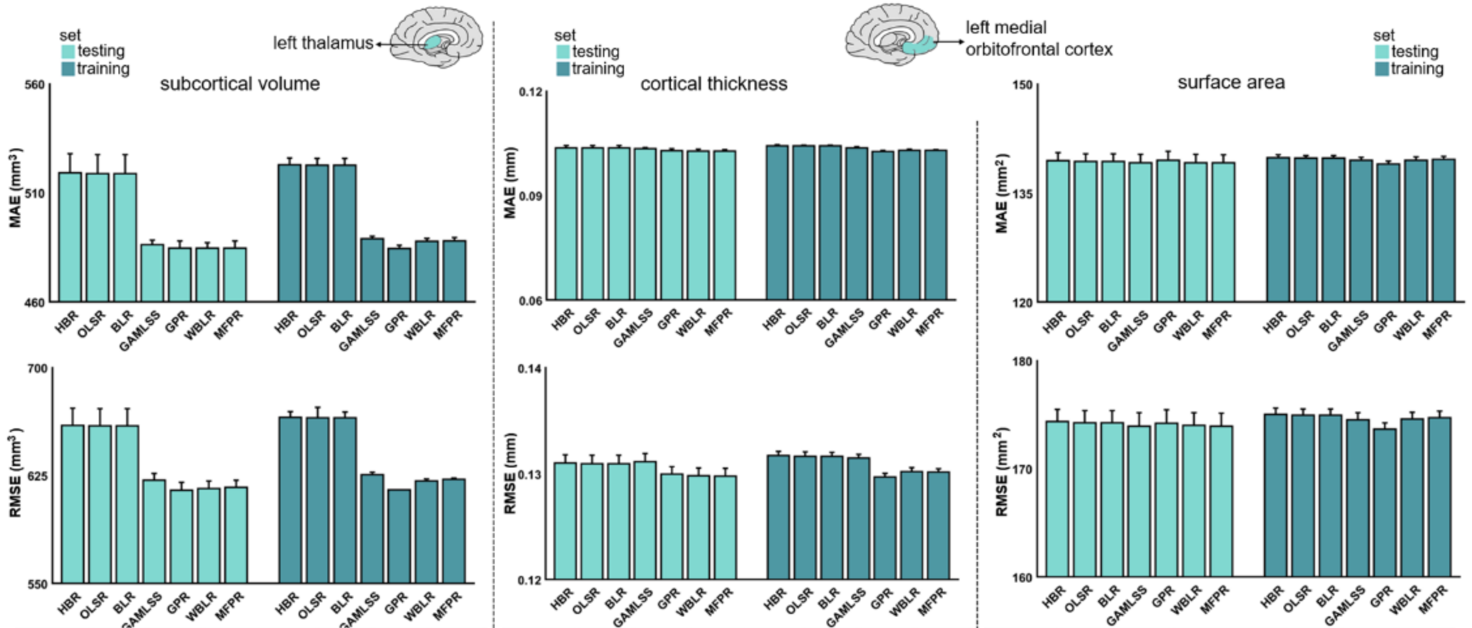


| algorithm | HBR | OLSR | BLR | GAMLSS | GPR | WBLR | MFPR |
|---|---|---|---|---|---|---|---|
| CPU time (seconds) | 904.84 ±524.52 | 0.01 ±0.001 | 5.35 ±0.35 | 3.89 ±0.18 | 1926.40 ±6.81 | 9.18 ±13.52 | 0.46 ±0.12 |

| algorithm | HBR | OLSR | BLR | GAMLSS | GPR | WBLR | MFPR |
|---|---|---|---|---|---|---|---|
| CPU time (seconds) | 921.68 ±139.45 | 0.01 ±0.001 | 4.29 ±0.02 | 3.42 ±0.41 | 1716.21 ±12.62 | 1.68 ±0.15 | 0.43 ±0.01 |

| algorithm | HBR | OLSR | BLR | GAMLSS | GPR | WBLR | MFPR |
|---|---|---|---|---|---|---|---|
| CPU time (seconds) | 298.16 ±55.38 | 0.01 ±0.001 | 4.44 ±0.07 | 3.57 ±0.17 | 1731.21 ±5.28 | 0.90 ±0.03 | 0.39 ±0.01 |

**Supplementary Figure S9. Illustrative examples of the comparative performance of HBR, OLSR, BLR, GAMLSS, GPR, WBLR, and MFPR-derived models in males.** Region-specific models with the optimized covariate combination were estimated in males and females separately using Hierarchical Bayesian Regression (HBR), Ordinary Least Squares Regression (OLSR), Bayesian Linear Regression (BLR), Generalized Additive Models for Location, Scale, and Shape (GAMLSS), Gaussian Process Regression (GPR), Warped Bayesian Linear Regression (WBLR), and Multivariable Fractional Polynomial Regression (MFPR). Model performance was assessed in terms of mean absolute error (MAE), root mean square error (RMSE), and central processing unit (CPU). The MAE, RMSE, and CPU time of the models for left thalamic volume (left panel), the left medial orbitofrontal cortical thickness (middle panel), and surface area (right panel) as exemplars are presented here for males and in figure 4 in the main text for females.

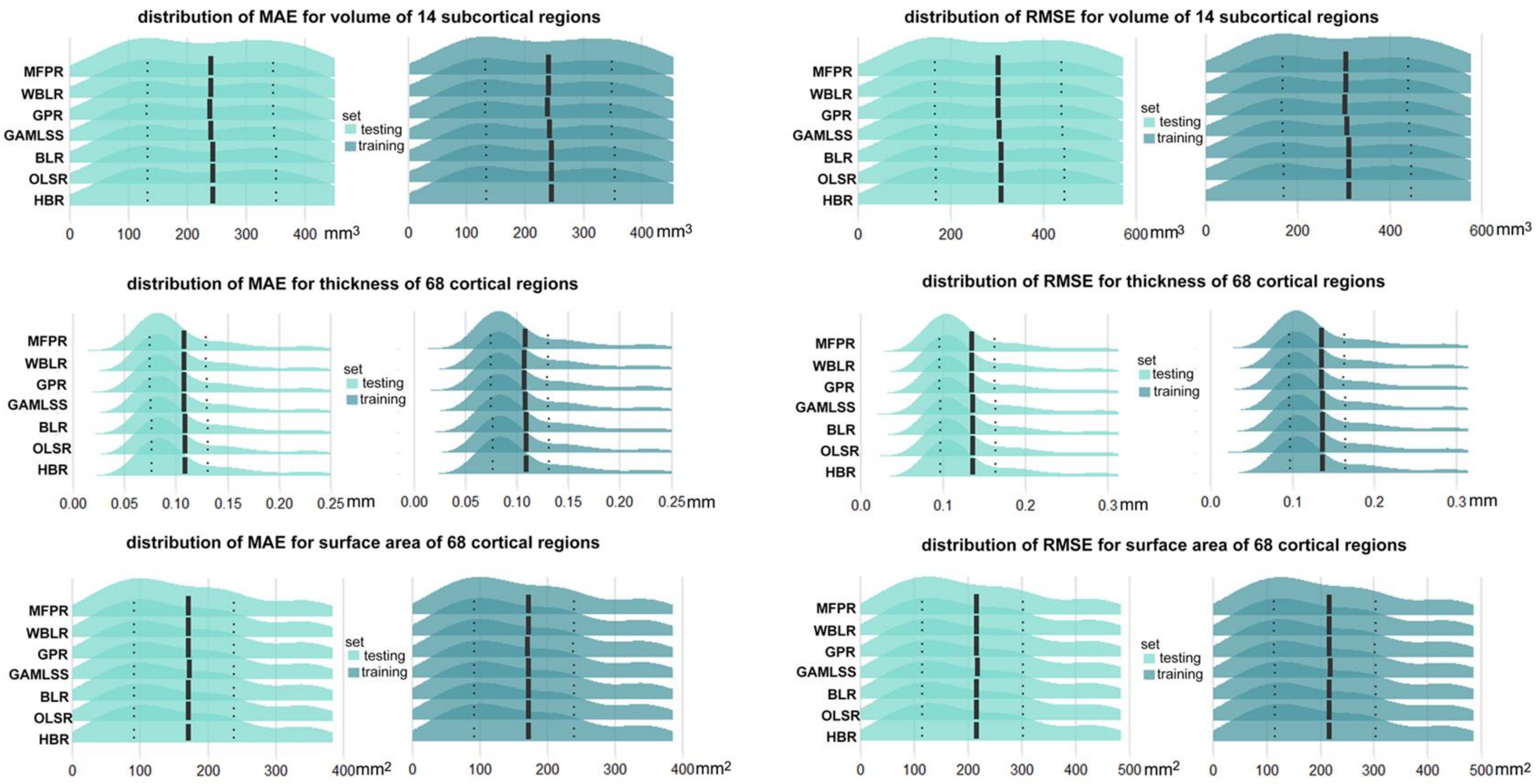


**Supplementary Figure S10. Comparative performance of all region-specific optimized HBR, OLSR, BLR, GAMLSS, GPR, HBR, WBLR, and MFPR-derived models in females.** The figure presents the distribution of the mean absolute error (MAE) and the root mean square error (RMSE) for all region-specific models derived from Hierarchical Bayesian Regression (HBR), Ordinary Least Squares Regression (OLSR), Bayesian Linear Regression (BLR), Generalized Additive Models for Location, Scale, and Shape (GAMLSS), Gaussian Process Regression (GPR), Warped Bayesian Linear Regression (WBLR), and Multivariable Fractional Polynomial Regression (MFPR). The mean MAE and RMSE values are marked with solid vertical lines, and the 25th percentiles and 75th percentiles are marked with dotted vertical lines.

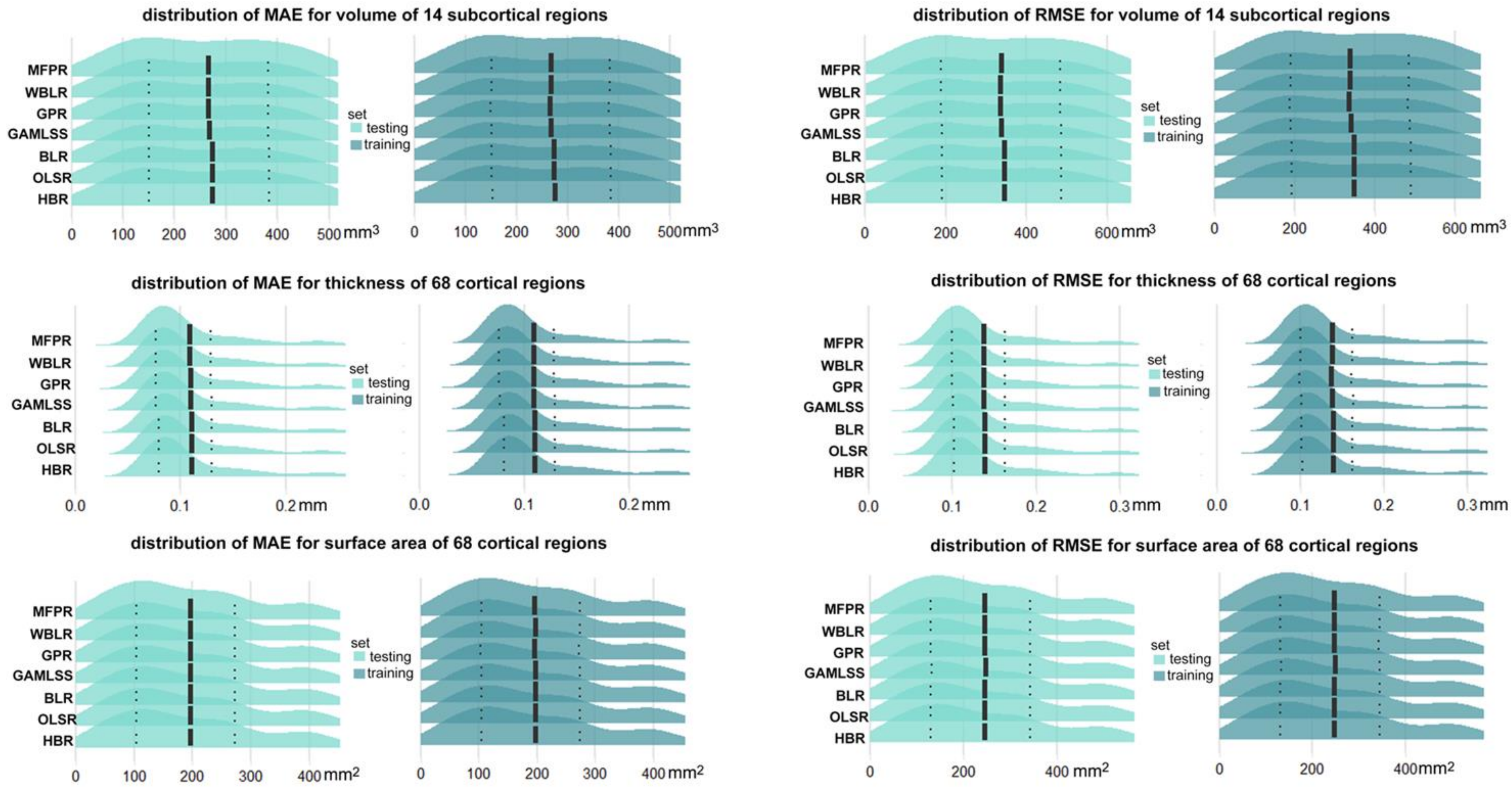


**Supplementary Figure S11. Comparative performance of all region-specific optimized HBR, OLSR, BLR, GAMLSS, GPR, HBR, WBLR, and MFPR-derived models in males**. The figure presents the distribution of the mean absolute error (MAE) and the root mean square error (RMSE) for all region-specific models derived from Hierarchical Bayesian Regression (HBR), Ordinary Least Squares Regression (OLSR), Bayesian Linear Regression (BLR), Generalized Additive Models for Location, Scale, and Shape (GAMLSS), Gaussian Process Regression (GPR), Warped Bayesian Linear Regression (WBLR), and Multivariable Fractional Polynomial Regression (MFPR). The mean MAE and RMSE values are marked with solid vertical lines, and the 25th percentiles and 75th percentiles are marked with dotted vertical lines.

**S5. Sensitivity analyses**
**S5.1 Sample size**

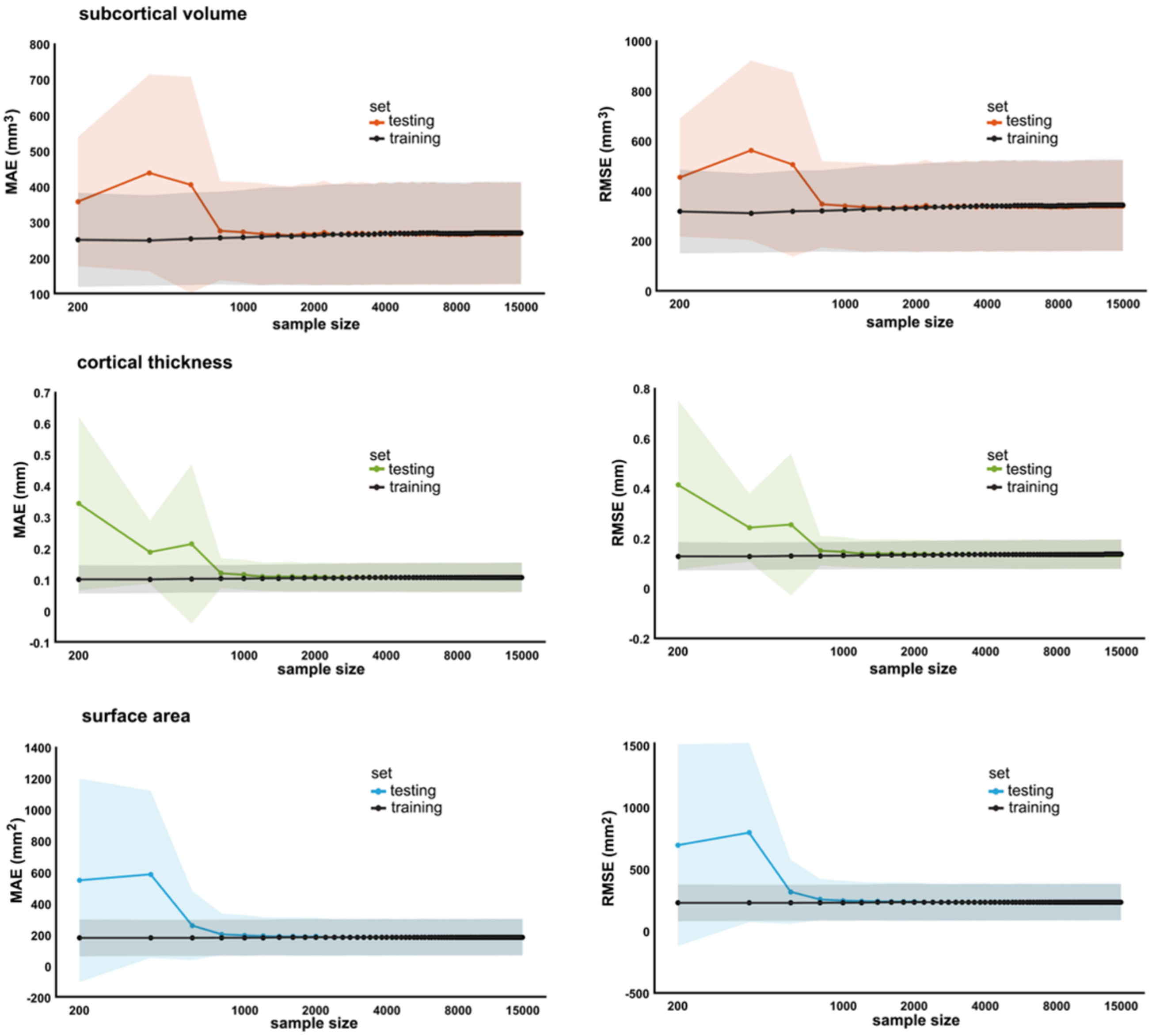


**Supplementary Figure S12. Performance of region-specific MFPR-derived models as a function of sample size for males.** Models for each regional morphometric measure were estimated in random sex-specific subsets of 200 to 15,000 participants, in increments of 200, generated from the study sample. Each line represents the values of the mean absolute error (MAE), or root mean square error (RMSE) derived from the optimized Multivariable Fractional Polynomial Regression (MFPR) models of each regional morphometric measure as a function of sample size. The pattern identified was identical in both sexes. The data for males are shown here and in figure 5 in the main text for females.

### S5.2 Age bins

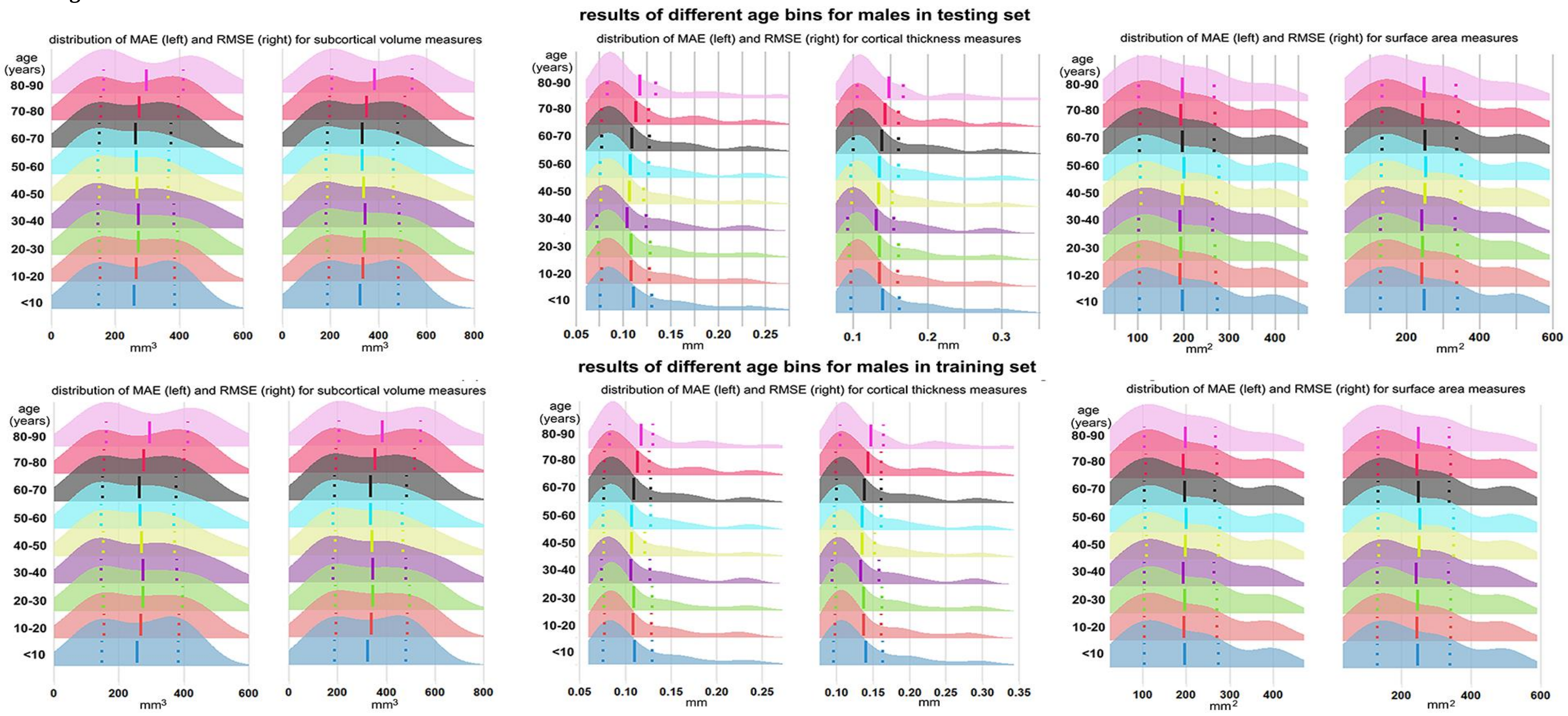


**Supplementary Figure S13. Performance of region-specific models in distinct age groups for males.** Sex- and region-specific models for all morphometric measures for different age groups were estimated by partitioning the sex-specific training and testing subsets of the study sample into nine age bins (i.e., age≤10 years; 10<age≤20 years; 20<age≤30 years; 30<age≤40 years; 40<age≤50 years; 50<age≤60 years; 60<age≤70 years; 70<age≤80 years; 80<age≤90 years). Details are provided in appendix 5. The pattern was identical in both sexes. Supplementary figure S9 presents the distribution of the mean absolute error (MAE) and the root mean square error (RMSE) across all region-specific models in males in the training (upper panel) and test subset (lower panel). The results for females are presented in figure 6 in the main text.

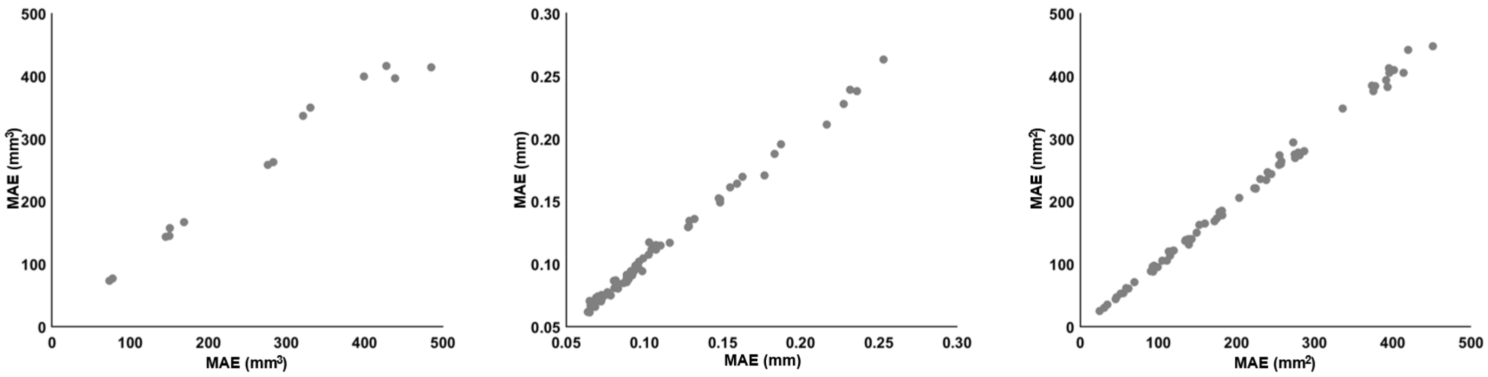


**Supplementary Figure S13 (continued). Scatter plots between the MAE values of the models within the first age bin (age⩽10 years) of male participants with those derived from the entire male sample.** Across all age bins, the correlation coefficient between the MAE or RMSE values of the sex- and region-specific models obtained from the full study sample and MAE or RMSE values of the corresponding models estimated in each age bin were all greater than 0·98. Same pattern was observed for all age bins of both sexes.

**S5. 3 Sensitivity analyses pertaining to the GAMLSS**

Prior studies have focused on GAMLSS algorithms and their relative performance in normative modeling of brain morphometric data. A key example is the paper by Dinga and colleagues.[12] As shown in supplementary figures S14 and S15 for females and males below, five models were compared: Model 1 is a Gaussian model with linear effect; Model 2 is a non-linear Gaussian homoscedastic model; Model 3 is a non-linear Gaussian heteroscedastic model; Model 4 is a non-linear heteroscedastic SHASH model; and Model 5 is a non-linear heteroscedastic SHASH model where location, scale, and shape also depend on age.

Model 1: regional neuromorphometric measure ~ $N(\mu, \sigma), \mu = \beta_0 + \beta_{age} * age$
Model 2: regional neuromorphometric measure ~ $N(\mu, \sigma), \mu = f_\mu(age)$
Model 3: regional neuromorphometric measure ~ $N(\mu, \sigma), \mu = f_\mu(age), \sigma = f_\sigma(age)$
Model 4: regional neuromorphometric measure ~ $SHASH(\mu, \sigma, \nu, \tau), \mu = f_\mu(age), \sigma = f_\sigma(age), \nu = \beta_{nu}, \tau = \beta_{tau}$
Model 5: regional neuromorphometric measure ~ $SHASH(\mu, \sigma, \nu, \tau), \mu = f_\mu(age), \sigma = f_\sigma(age), \nu = f_\nu(age),\ \tau = f_\tau(age)$

Model 4 is the model used in our manuscript. All other models are from Dinga et al.[12] These five GAMLSS models were compared in terms of MAE (mean absolute error), kurtosis, and skewness. All models have similar MAE, while model 4 and model 5 performed best in terms of kurtosis and skewness (i.e., for kurtosis and skewness, closer to 0 is better). Since differences in the performance of models 4 and 5 are negligible, the results support our choice of using model 4, which is also simpler (figure 2 in the main text). The same pattern was found for RMSE (root mean squared error) and EV (explained variance).

We also compared the performance of the GAMLSS models using either the "caret" or "gamlss" packages in terms of MAE, kurtosis, and skewness using the same sample as above. The results shown in supplementary figures S14-S15 demonstrate the similarity between the two packages in MAE. The output of the "caret" package used in our paper performed better in terms of the model fit of kurtosis and skewness (for kurtosis and skewness, closer to 0 is better) for the testing set. These analyses all support the choice of the package used in our manuscript.

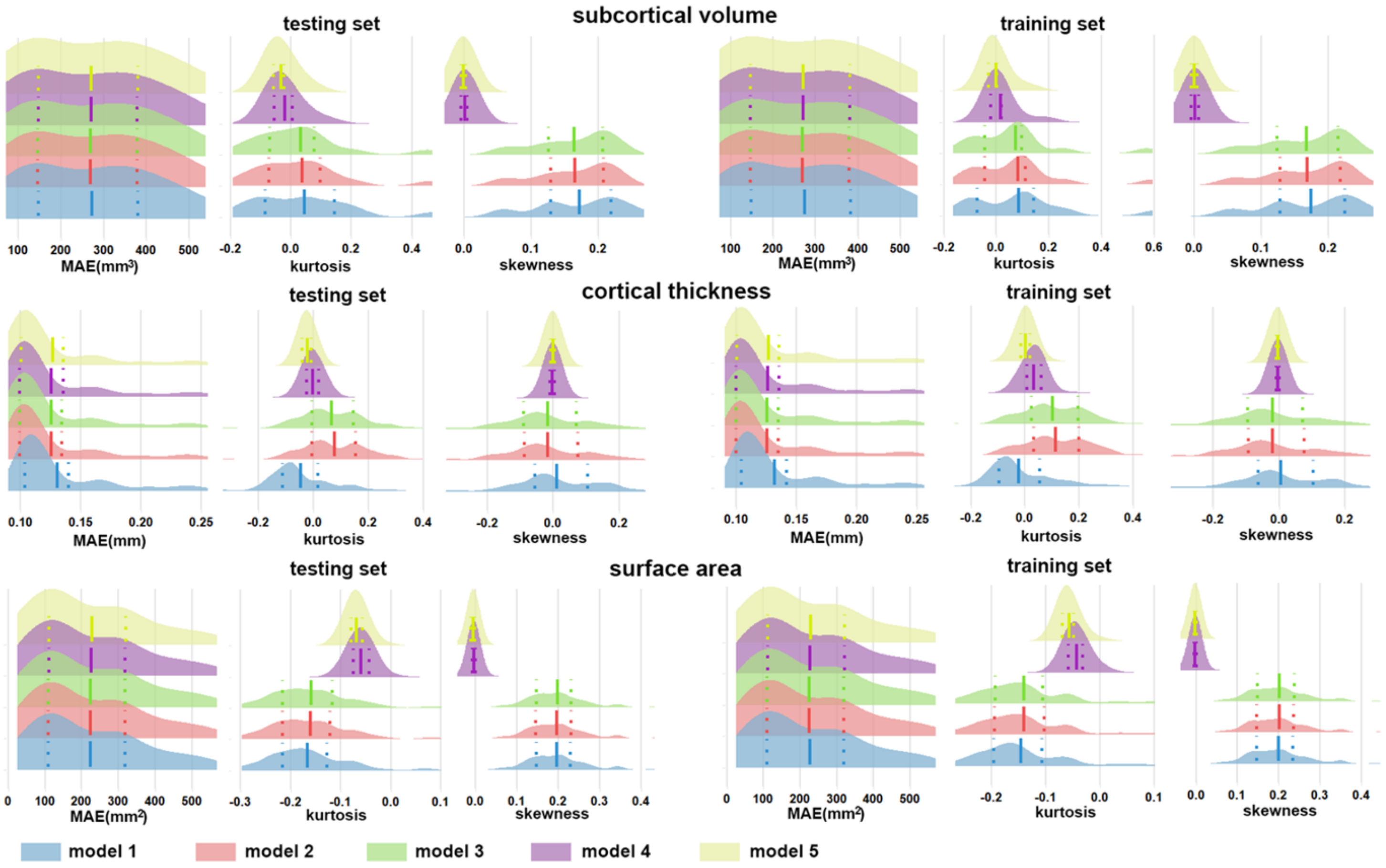


**Supplementary Figure S14**. **Comparative evaluations of five GAMLSS models in females**. The models were compared in terms of MAE (mean absolute error), kurtosis, and skewness. The model used in our paper is model 4, all other models are from Dinga et al.[12] All models have similar MAE, while model 4 and model 5 performed best in terms of kurtosis and skewness (i.e., for kurtosis and skewness, closer to 0 is better). Since differences in the performance of models 4 and 5 are negligible, the results support our choice of using model 4.

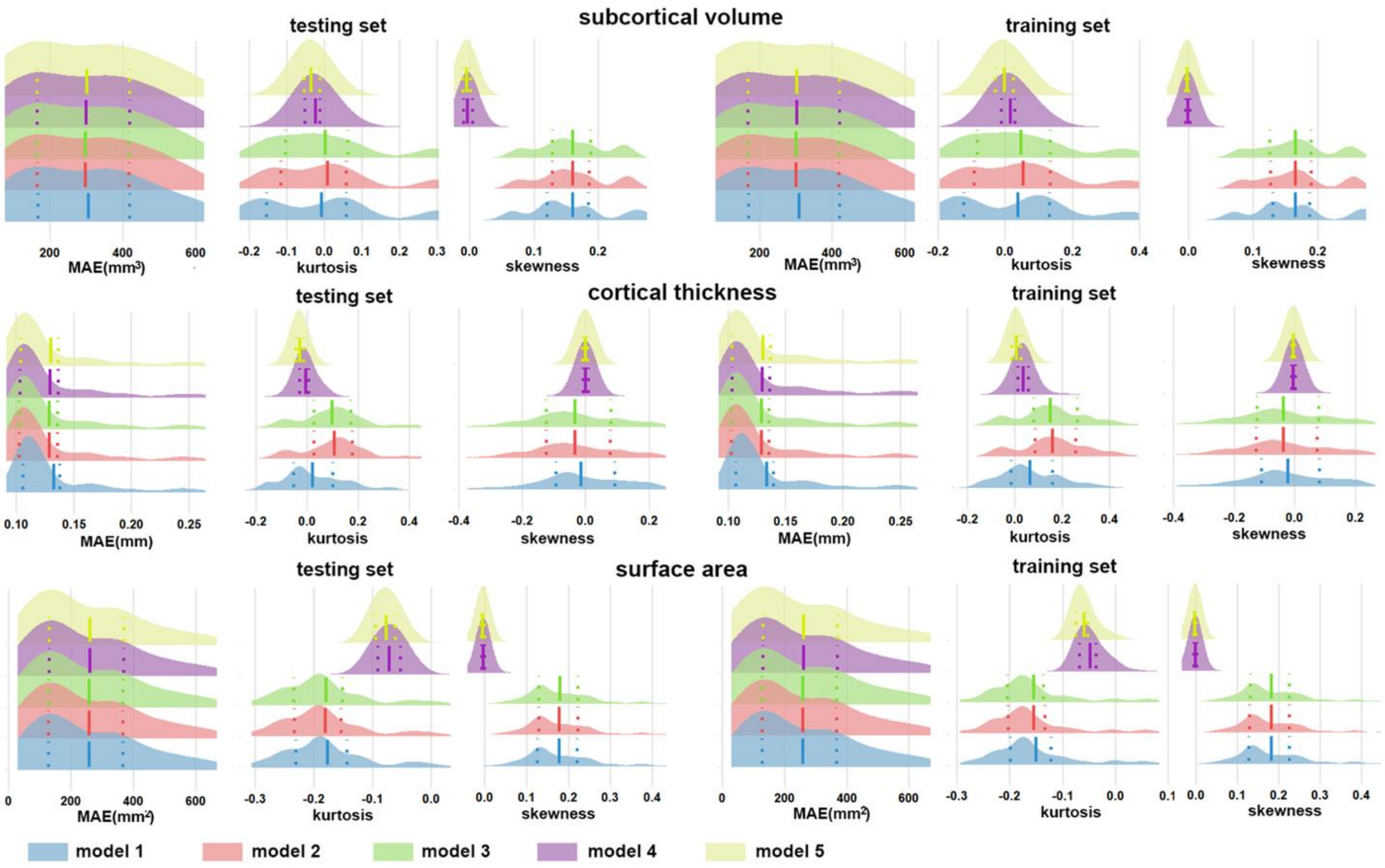


**Supplementary Figure S15**. **Comparative evaluations of five GAMLSS models in males**. The models were compared in terms of MAE (mean absolute error), kurtosis, and skewness. The model used in our paper is model 4, all other models are from Dinga et al.[12] All models have similar MAE, while model 4 and model 5 performed best in terms of kurtosis and skewness (i.e., for kurtosis and skewness, closer to 0 is better). Since differences in the performance of models 4 and 5 are negligible, the results support our choice of using model 4.

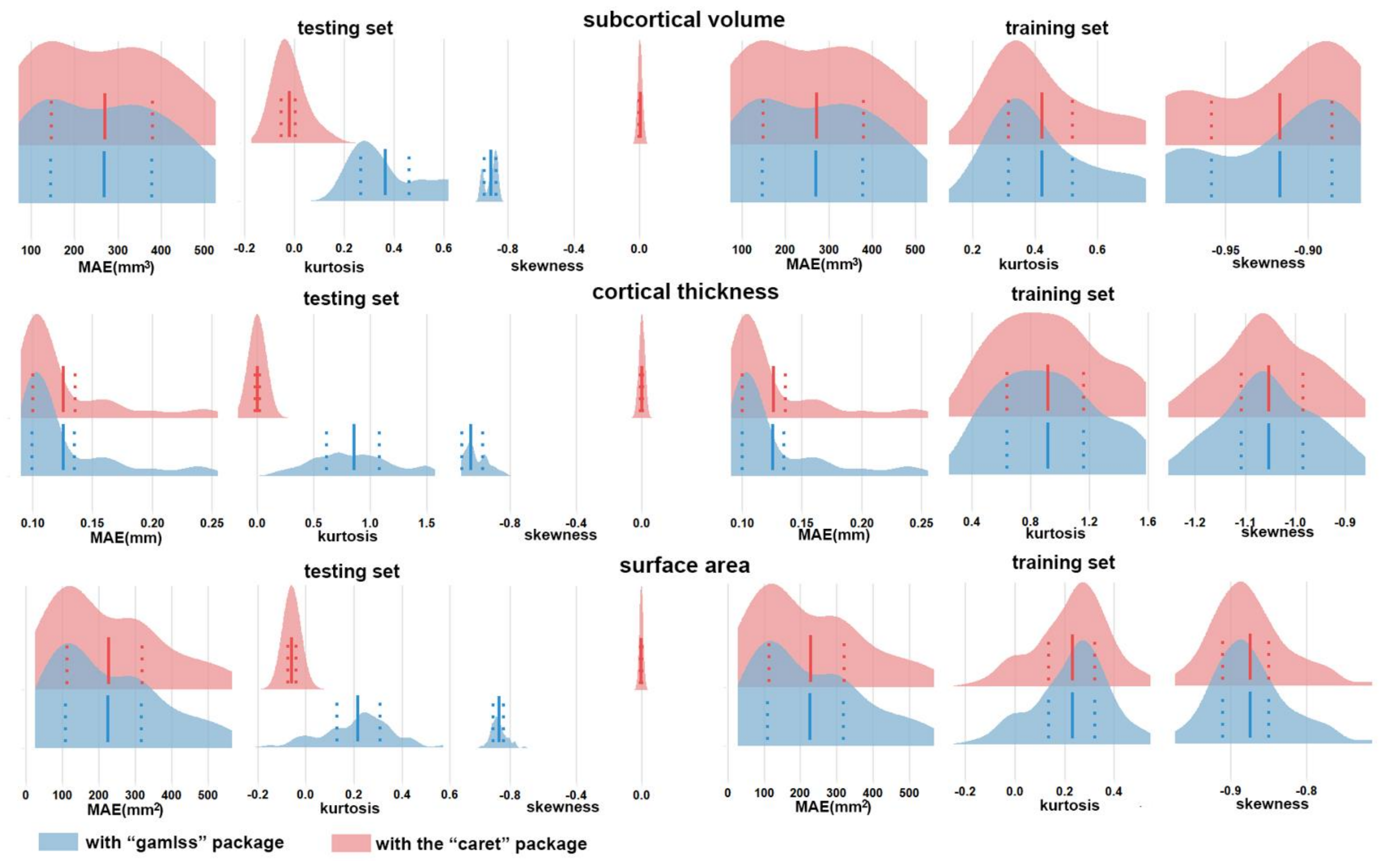


**Supplementary Figure S16. Comparative evaluation of GAMLSS models using the "caret" and "gamlss" software in females.** GAMLSS models using the "caret" (pink) and "gamlss" (blue) packages in terms of MAE (mean absolute error), kurtosis, and skewness. Model performance in terms of MAE was similar for both packages. The output of the "caret" package used in our paper performed better in terms of the model fit of kurtosis and skewness (for kurtosis and skewness, closer to 0 is better) for the testing set.

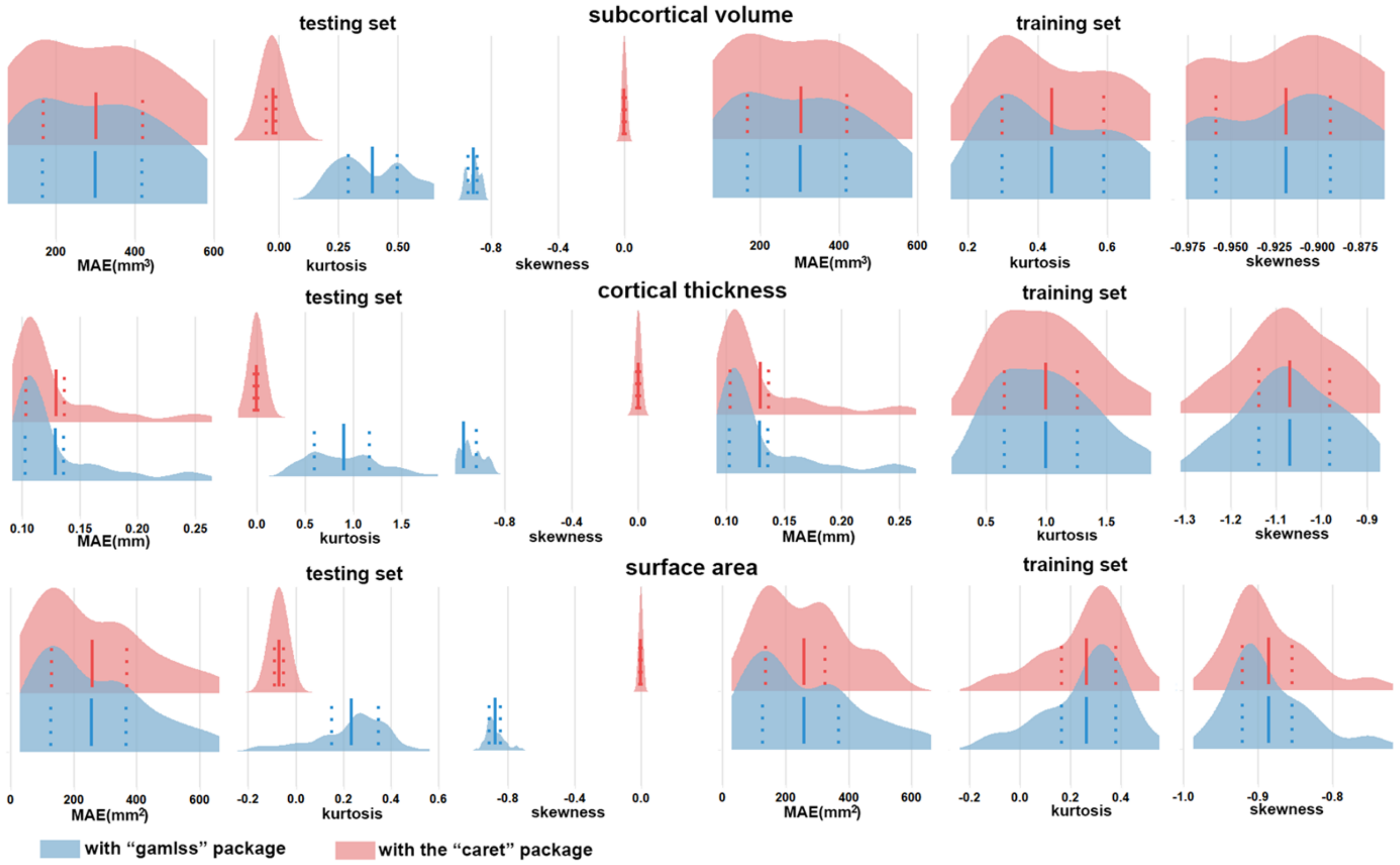


**Supplementary Figure S17. Comparative evaluation of GAMLSS models using the “caret” and “gamlss” software in males.** GAMLSS models using the “caret” (pink) and “gamlss” (blue) packages in terms of MAE (mean absolute error), kurtosis, and skewness. Model performance in terms of MAE was similar for both packages. The output of the “caret” package used in our paper performed better in terms of the model fit of kurtosis and skewness (for kurtosis and skewness, closer to 0 is better) for the testing set.

### S5.4 Sensitivity analyses pertaining to site

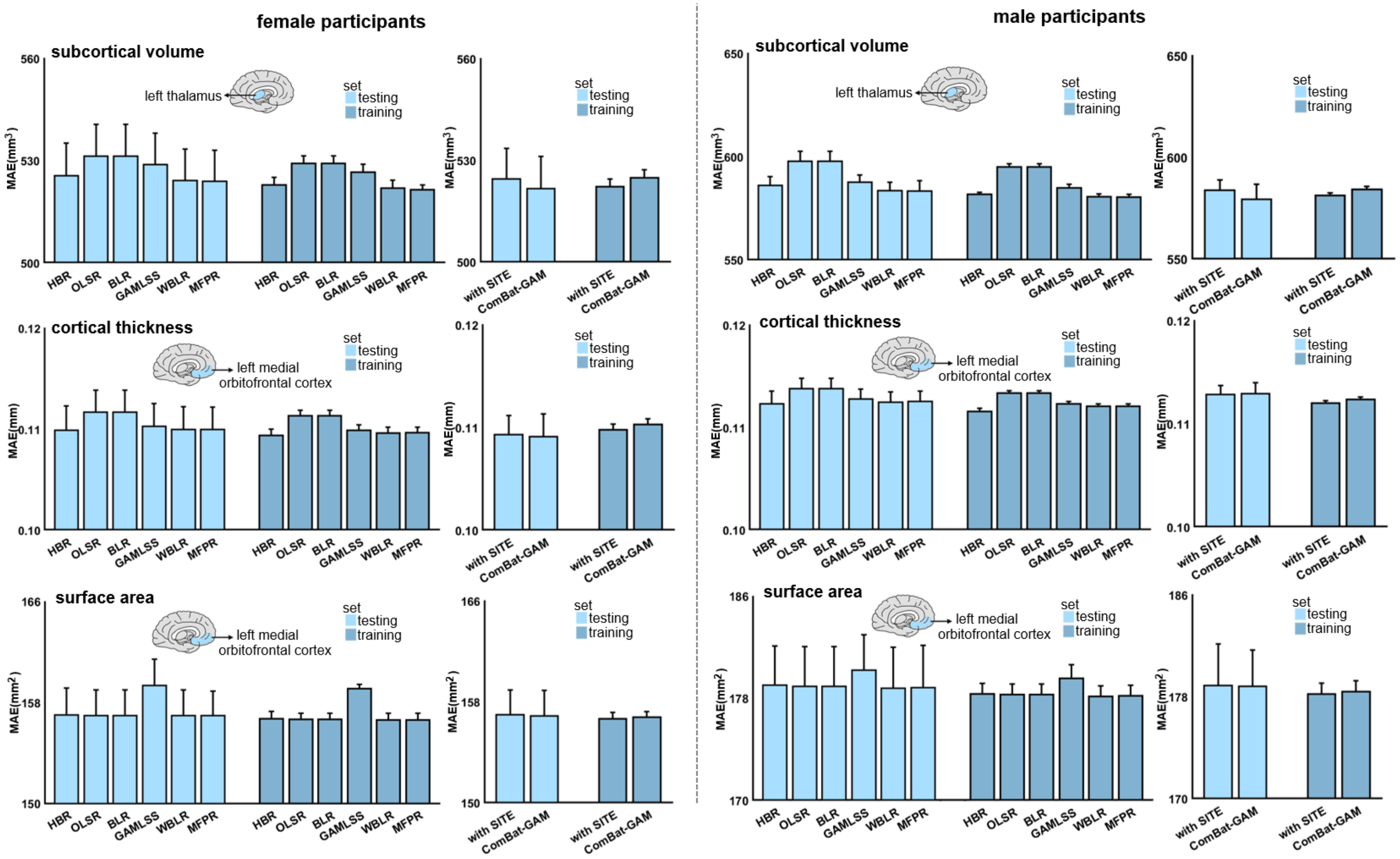


**Supplementary Figure S18. Effect of site handling on algorithm performance.** The performance of algorithms [Hierarchical Bayesian Regression (HBR); Ordinary Least Squares Regression (OLSR); Bayesian Linear Regression (BLR); Generalized Additive Models for Location, Scale, and Shape (GAMLSS); Warped Bayesian Linear Regression (WBLR); Multivariable Fractional Polynomial Regression (MFPR)] was compared when site was modeled either as a random factor or removed by site harmonisation using ComBat-GAM. The top performing algorithm when the site was used as a random effect was still the MFPR, followed closely by WBLR. Further, the model performance of the MFPR algorithm in terms of MAE was similar regardless of how site was handled.

**S6. Clinical relevance of normative brain modeling for mental illness**

We tested the relative advantage of the brain regional normative deviation scores (Z-scores) compared to observed neuromorphometric data for the discrimination of healthy individuals from those with early onset psychosis and for the prediction of psychotic symptom severity. To achieve this, we used data from the Human Connectome Project (HCP) Early Psychosis Study which comprises cross-sectional neuroimaging data from 91 individuals with early psychosis and 57 healthy individuals (total sample: 48 females/100 males; age range 16.67-35.67 years). T1-weighted images were downloaded from the HCP repository (https://www.humanconnectome.org/study/human-connectome-project-for-early-psychosis) and preprocessed using FreeSurfer (version 7.1.0) to generate regional brain morphometric data. Then brain regional Z-scores were derived from each of the algorithms examined here. As described in the main manuscript, the classification accuracy of support vector classifiers (SVC) using brain regional Z-scores were compared to that of the classifier using observed data. Predictive accuracy for psychotic symptom severity was assessed using ridge regression models using brain regional Z-scores compared to that of the classifier using observed data.

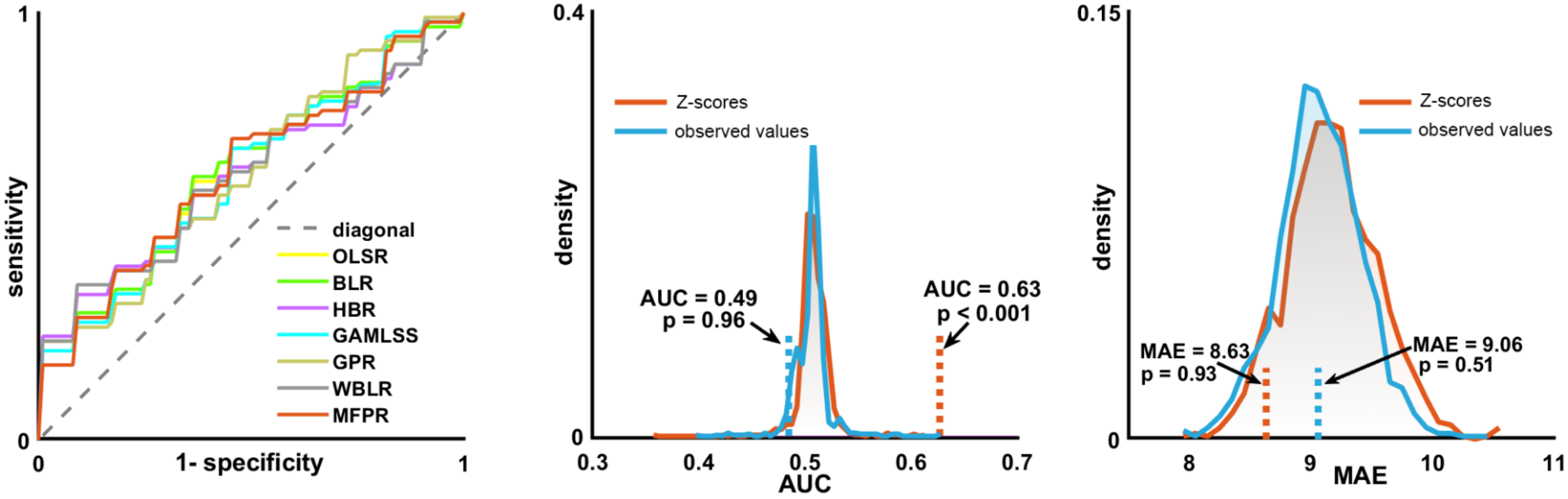


**Supplementary Figure S19. Comparative accuracy of diagnostic classification and symptom severity prediction in psychosis using either brain regional Z-scores or observed neuromorphometric data of the Human Connectome Project-Early Psychosis cohort.** Left panel: Receiver Operating Characteristic Curves (ROC) of support vector classifiers (SCV) using brain regional Z-scores from each of the algorithms examined here. Middle Panel: Comparative performance in accuracy of diagnostic classification of the SVC using either Z-scores (red) or observed data (blue). Right panel: Predictive accuracy for psychotic symptom severity of ridge regression models using either Z-scores (red) or observed data (blue). In both these panels, two null distributions from the permutation tests are presented: one that illustrates the performance of models using the observed data (distributions with blue outlines) and the other illustrating the performance of models using Z-scores (distributions with red outlines). AUC=area under the curve; MAE=mean absolute error; OLSR=Ordinary Least Squares Regression; BLR=Bayesian Linear Regression; HBR=Hierarchical Bayesian Regression; GAMLSS=Generalized Additive Models for Location, Scale, and Shape; GPR=Gaussian Process Regression; WBLR=Warped Bayesian Linear Regression; MFPR=Multivariable Fractional Polynomial Regression.

**S7. List of appendices 2-6 in Excel format**

Appendix 2: Supplementary Table S1. Datasets included in the model optimization sample with access information.
Appendix 3: Supplementary Table S2. Performance of the eight algorithms for regional subcortical volume models in females
Appendix 3: Supplementary Table S3. Performance of the eight algorithms for regional subcortical volume models in males
Appendix 3: Supplementary Table S4. Performance of the eight algorithms for regional cortical thickness models in females
Appendix 3: Supplementary Table S5. Performance of the eight algorithms for regional cortical thickness models in males
Appendix 3: Supplementary Table S6. Performance of the eight algorithms for regional cortical surface area models in females
Appendix 3: Supplementary Table S7. Performance of the eight algorithms for regional cortical surface area models in males
Appendix 4: Supplementary Table S8. Performance of MFPR-derived models of regional subcortical volumes in females as a function of explanatory variables
Appendix 4: Supplementary Table S9. Performance of MFPR-derived models of regional subcortical volumes in males as a function of explanatory variables
Appendix 4: Supplementary Table S10. Performance of MFPR-derived models of regional cortical thickness in females as a function of explanatory variables
Appendix 4: Supplementary Table S11. Performance of MFPR-derived models of regional cortical thickness in males as a function of explanatory variables
Appendix 4: Supplementary Table S12. Performance of MFPR-derived models of regional cortical surface area in females as a function of explanatory variables
Appendix 4: Supplementary Table S13. Performance of MFPR-derived models of regional cortical surface area in males as a function of explanatory variables
Appendix 5: Supplementary Table S14. Performance of MFPR-derived models of regional subcortical volumes per age bin in females
Appendix 5: Supplementary Table S15. Performance of MFPR-derived models of regional subcortical volumes per age bin in males
Appendix 5: Supplementary Table S16. Performance of MFPR-derived models of regional cortical thickness per age bin in females
Appendix 5: Supplementary Table S17. Performance of MFPR-derived models of regional cortical thickness per age bin in males
Appendix 5: Supplementary Table S18. Performance of MFPR-derived models of regional cortical surface area per age bin in females
Appendix 5: Supplementary Table S19. Performance of MFPR-derived models of regional cortical surface area per age bin in males
Appendix 6: Supplementary Table S20. Comparative performance and CPU time of optimized OLSR, BLR, HBR, GPR, GAMLSS, WBLR, and MFPR-derived models for regional subcortical volumes in females
Appendix 6: Supplementary Table S21. Comparative performance and CPU time of optimized OLSR, BLR, HBR, GPR, GAMLSS, WBLR, and MFPR-derived models for regional subcortical volumes in males
Appendix 6: Supplementary Table S22. Comparative performance and CPU time of optimized OLSR, BLR, HBR, GPR, GAMLSS, WBLR, and MFPR-derived models for regional cortical thickness in females
Appendix 6: Supplementary Table S23. Comparative performance and CPU time of optimized OLSR, BLR, HBR, GPR, GAMLSS, WBLR, and MFPR-derived models for regional cortical thickness in males
Appendix 6: Supplementary Table S24. Comparative performance and CPU time of optimized OLSR, BLR, HBR, GPR, GAMLSS, WBLR, and MFPR-derived models for regional cortical surface area in females
Appendix 6: Supplementary Table S25. Comparative performance and CPU time of optimized OLSR, BLR, HBR, GPR, GAMLSS, WBLR, and MFPR-derived models for regional cortical surface area in males
